# Design and Assembly of Combinatorial DNA Barcodes for Probe-based Genomics Applications

**DOI:** 10.64898/2026.08.22.746475

**Authors:** Zach Goode, Emily Tiedemann, Lamya Ben Ameur, Kenny Pavan, Karl Young, Melissa Sek, Alexander Nevue, Jinjin Zhu, Jessica Houghton, Ye Fu, Heike Boisvert, Arpiar Saunders

## Abstract

Probe-based genomics technologies are extending molecular analysis into intact tissues and fixed cells, yet strategies to decode complex experimental conditions encoded in cellular RNA remain limited. Here we present a modular framework that integrates custom software tools with purpose-built cloning reagents to design, assemble, validate, and deploy “combinatorial” DNA barcodes. Combinatorial barcodes comprise spatially adjacent collections of known sequences, enabling millions of unique molecules to be efficiently distinguished using a limited set of probes. Our software tools integrate with optimized assembly plasmids and whole plasmid long-read sequencing for high-fidelity construction and structural validation of diverse combinatorial barcode architectures. Assembled barcode libraries are flexibly transferred into user-modified expression vectors to support diverse downstream experimental applications. We showcase the versatility of this framework by assembling two structurally distinct combinatorial barcode libraries, each containing millions of unique sequences. Following rabies virus-based delivery to the mouse brain, we validate *in vivo* decoding of a combinatorial barcode architecture capable of distinguishing ∼16.3 million expressed RNAs through probe-based *in situ* sequencing. Our framework for flexible and accurate combinatorial barcode construction fills a technically demanding niche delivering cost-effective molecular reagents for multiplexed experimentation on current and evolving probe-based genomics platforms.

**GRAPHICAL ABSTRACT:** 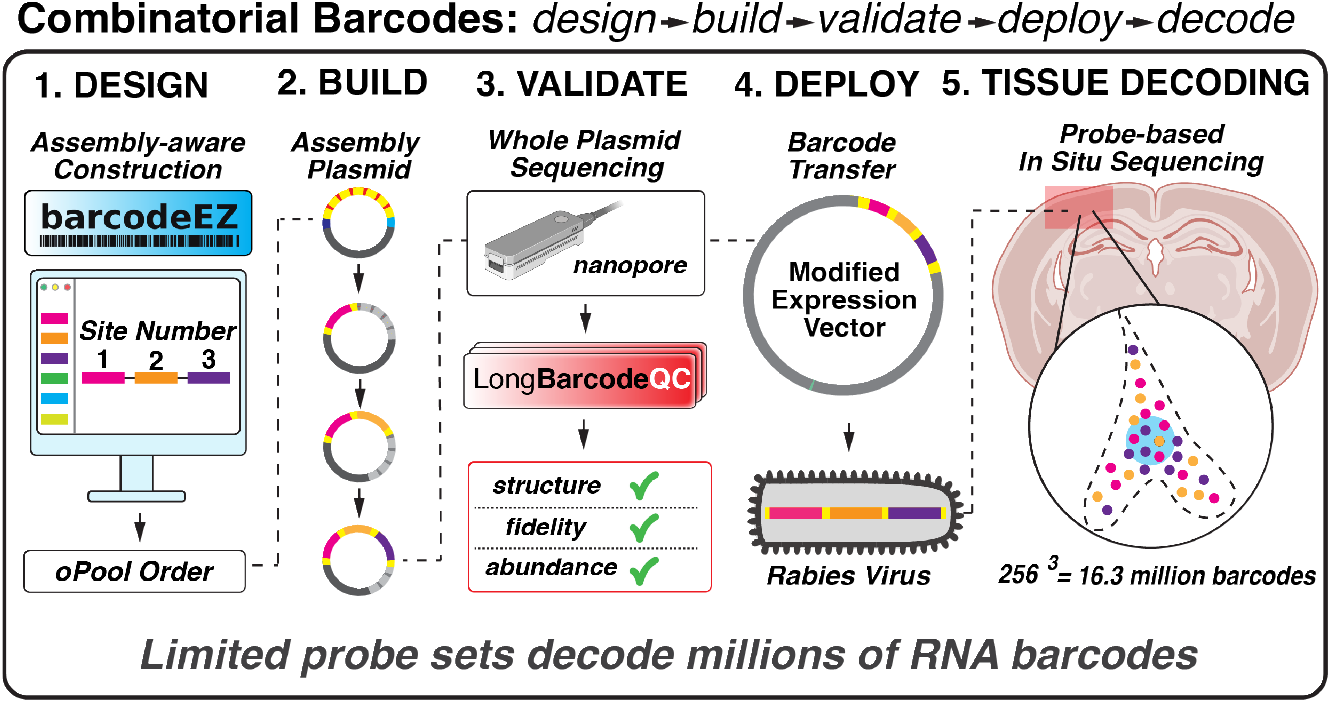

## INTRODUCTION

Since pioneering early demonstrations (1), the use of DNA and RNA sequences as “barcodes” to track experimental conditions has rapidly evolved into a central strategy for single-cell and functional genomics. In most implementations, short synthetic barcode sequences are generated through oligonucleotide synthesis and decoded alongside genome-wide RNA or DNA molecules using high-throughput flow cell–based sequencing. The accuracy and throughput of flow cell-based sequencing are well-suited to identify short barcode sequences from large sets, enabling millions of experimental conditions to be tracked in parallel. In neuroscience research alone, compact barcoding schemes coupled with single-cell RNA library generation and flow cell sequencing have enabled pooled CRISPR-based perturbation screens (2), cell type gene enhancer characterization (3), developmental lineage tracing (4–6), viral tropism analysis (7–9), and the mapping of complex anatomy, including neuronal projection systems (10–12) and synaptic networks (13–15).

In recent years, probe-based technologies -including multiplexed fluorescent *in situ* hybridization (FISH)(16, 17), *in situ* sequencing (18, 19), and fixed-sample transcriptome profiling (19) - are extending genome-scale molecular analyses into new experimental domains, including intact tissue and fixed and archival specimens (19, 20). For example, probe-based genomics methods enable analysis of molecularly defined cell populations in intact brain tissue, helping generate new insights into neural circuit structure-function relationships (21). In combination with straightforward cross-animal comparisons, such probe-based approaches have helped identify principles of brain development (22), aging (23, 24), and neurodegeneration (25, 26). One attractive feature of molecular analysis within intact tissue is the potential for more comprehensive sampling of experimentally relevant cells, which are otherwise subsampled by single-cell genomics approaches requiring tissue dissociation and microfluidic library generation. Recent advances are pushing intact tissue analysis from thin 10-20 μm sections to thicker 100 μm slabs, supporting volumetric reconstructions of millions of cells with retained anatomy (27). While probe-based approaches have begun to support high-throughput CRISPR screens in fixed cells (28–30), most current datasets are descriptive, involve cross-tissue comparisons, or track only a small number of experimental conditions per experiment (31, 32). Enabling highly multiplexed experimentation on current and future probe-based genomics platforms requires methods that dovetail with platform-specific chemistry to accurately discern millions of experimental conditions encoded in the RNA of single cells. Thus barcoding schemes that use limited probe sets to decode millions of experimentally linked RNA or DNA molecules remain a critical experimental need.

One powerful strategy for scaling the number of unique sequences detectable with probes is through a combinatorial barcode design. In this framework, each barcode comprises multiple sites containing user-defined sequences. The number of combinatorial barcodes grows exponentially as a function of the number of sites and site-specific sequences, enabling tens to hundreds of site-specific probes to decode millions of unique sequences. Combinatorial barcodes are gaining traction: for example, recent efforts have used probe-based combinatorial barcode decoding to CRISPR manipulations (28, 33) or developmental lineages of single cells (34, 35) or. While advances in synthesis of pooled oligonucleotides enable cost-effective generation of short synthetic DNA fragments, direct synthesis of full-length combinatorial barcode libraries remains prohibitively expensive at scale. Therefore, the most straightforward way to flexibly construct combinatorial barcode libraries is through serial assembly of shorter synthetic sequences. However, high-fidelity serial restriction cloning remains a major challenge, as cloning errors and unwanted plasmid byproducts accumulate during site-wise assembly, issues that compound with more complexity combinatorial barcode architectures.

To overcome these limitations, we present an integrated design–assemble–validate–deploy framework for constructing combinatorial barcode libraries based on custom cloning reagents, software tools, and nanopore sequencing (**Figure 1**). We introduce streamlined Assembly Plasmids (AP) with a six-site cloning architecture that separates serial assembly from downstream use in expression vectors, enabling optimized assembly followed by modular experimental porting. Our AP design extends the elegant restriction site “trio” strategy introduced by the single-cell-RNA-sequencing-compatible tracer for identifying clonal relationships (STICR) technology for site-wise barcode cloning and removal of insertion failures (5). To support flexible generation of distinct combinatorial barcode architectures, we developed *BarcodeEZ*, a Python-based software package that implements assembly-aware combinatorial barcode design, and *LongBarcodeQC*, a complementary command-line tool that processes whole plasmid long-read sequencing data and provides detailed secondary analysis of barcode composition and integrity. Combinatorial barcode libraries completed in the AP are introduced into modified expression vectors using a simple and modular cloning strategy. We illustrate the flexibility and accuracy of this system by creating “TritSeq” and “PadlockSeq” plasmid libraries, distinct combinatorial barcode designs that encode millions of unique sequences amenable to probe-based readout. We demonstrate *in vivo* decoding of the PadlockSeq design through probe-based *in situ* sequencing after viral expression in the mouse brain, validating the PadlockSeq barcode architecture as suitable for sensitive and accurate tracking millions of unique sequences encoded in cellular RNA. By combining efficient assembly, long-read quality control, and modular expression vector deployment, our platform enables complex combinatorial barcode libraries to be designed, assembled, and benchmarked with ease, facilitating the generation and reuse of high-quality reagents for multiplexed experimentation on diverse and evolving probe-based genomics platforms. The Assembly Plasmids, software tools, and constructed libraries are made publicly available to support community use and adoption.

**Figure 1.**
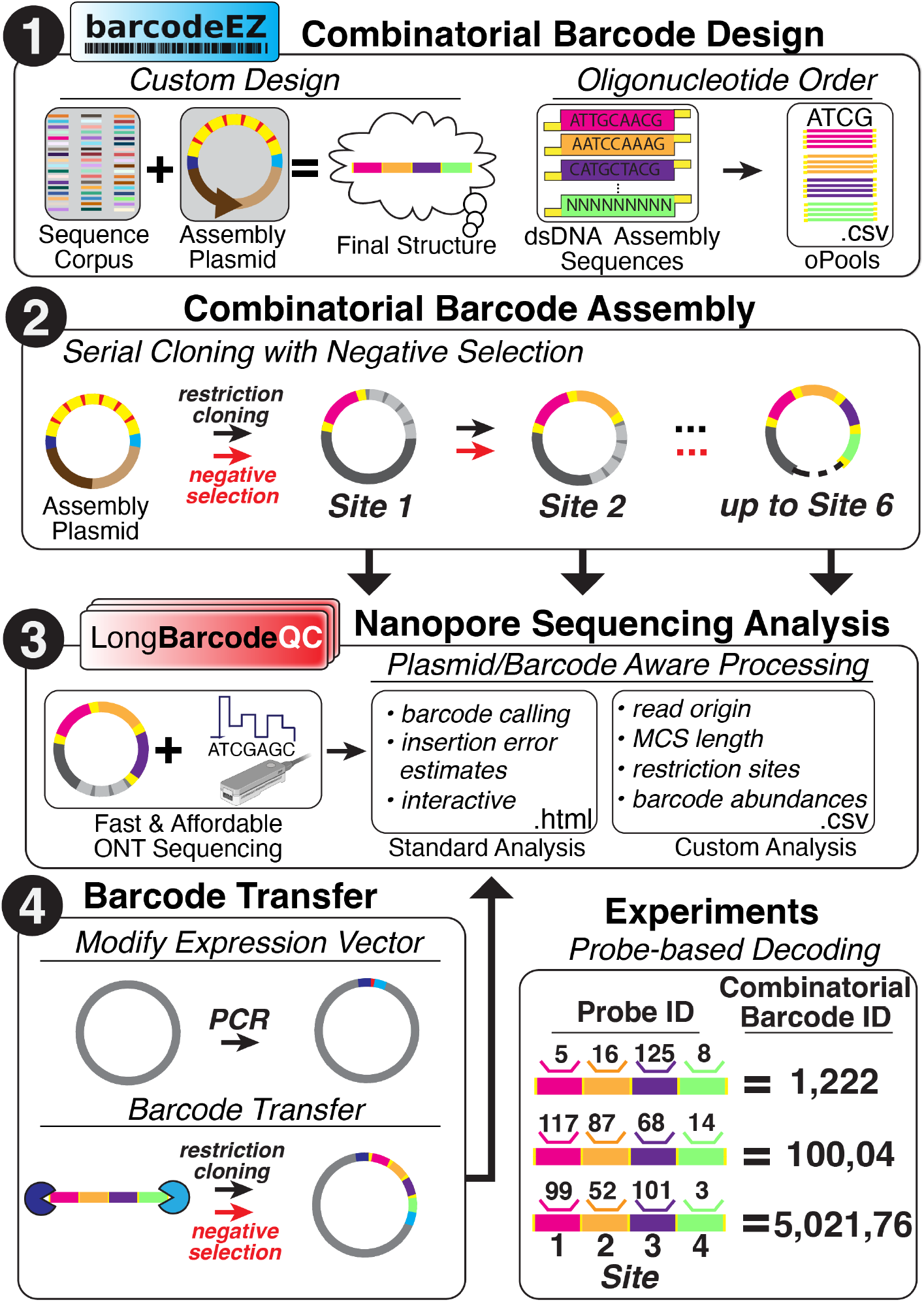
Overview of the integrated molecular and computational framework for combinatorial barcode design, assembly and quality control. Ordered schematic of the major molecular and computational components of the described framework. 1) Combinatorial barcode structures are designed in *BarcodeEZ*, an object-oriented pip-installable Python package. Designs are assembled from 60-mer synthetic sequence corpora orthogonal to model-species transcriptomes and modified for site-specific cloning into the Assembly Plasmid (AP). Flexible filters ensure seamless assembly and avoid sequences with undesired user-defined properties. *BarcodeEZ* exports designed barcodes as single-stranded DNA oligonucleotides for pooled commercial synthesis. 2) Combinatorial barcodes are cloned into a custom-designed AP using serial site-specific restriction cloning and negative selection. The AP has a multiple cloning site (MCS) containing six restriction site trios and a streamlined design to avoid unwanted cloning byproducts. 3) Long-read nanopore sequencing is used to assess site-specific barcode assembly with *LongBarcodeQC*, a pip-installable Python package. *LongBarcodeQC* uses expected barcode and plasmid sequences (with the AP as a default), supporting rapid and affordable quality control of the barcoded MCS and diversity characterization through a standard interactive HTML report and verbose outputs for custom downstream analysis. 4) Fully assembled combinatorial barcode sequences are transferred from the finalized AP library into user-defined expression vectors (EV) through a two-stage process. First, polymerase chain reaction (PCR) is used for targeted modification of EVs with restriction sites for barcode transfer cloning and negative selection. Second, restriction “Transfer” sites flanking the barcoded MCS are used to digest the AP and EV, generating compatible sticky-ended fragments for subsequent ligation and negative selection to remove non-barcoded EV. Orthogonal AP/EV antibiotic resistance limits residual AP carryover. *LongBarcodeQC* analysis of barcoded EV supports final quality control of barcode transfer. 5) Once experimentally deployed in the EV, each combinatorial barcode is decoded using probes targeting its site-specific sequence components.

## MATERIALS AND METHODS

### Software Development

*BarcodeEZ: a Python package for combinatorial barcode design. BarcodeEZ* is a Python (developed under Python 3.10; tested on Python 3.10 and 3.14) package developed to generate orthogonal combinatorial DNA barcode libraries for plasmid-based assembly. The software is implemented as an importable Python library. *BarcodeEZ* (v1.0.0) is freely available under the MIT license at https://github.com/ArpiarSaundersLab/barcodeEZ and can be installed with pip install barcodeEZ==1.0.0. Documentation: https://arpiarsaunderslab.github.io/barcodeEZ/.

*BarcodeEZ* constructs barcode libraries from a precomputed corpus of synthetic DNA sequences generated under defined sequence constraints. The default corpus comprises ∼30, 000 unique 60-mer sequences with GC content constrained to 41–59%, no homopolymer runs exceeding three bases, nearest-neighbor Levenshtein edit distance > 18, and exclusion of restriction enzyme recognition sites required for Assembly Plasmid cloning and transfer. To promote biological orthogonality, candidate sequences were screened using BLASTn against RefSeq-annotated transcriptomes of human (GRCh38.p14; GCF_000001405.40), mouse (GRCm39; GCF_000001635.27), and rhesus macaque (Mmul_10; GCF_003339765.1), downloaded from NCBI “ rna_from_genomic ” FASTA files. Sequences exhibiting alignments ≥18 nucleotides at 100% identity or ≥25 nucleotides at ≥85% identity or identical to any transcript were excluded from the corpus.

Barcode designs are instantiated as multi-site objects (`Barcodes()`), in which users specify the number of barcode insertion sites and, optionally, the number of internal positions per site. Barcode sequences are generated for each site using `generate_barcodes()` by sampling from the sequence corpus. The `validate()` method enforces sequence constraints by excluding candidate barcodes that introduce disallowed restriction sites or user-defined motifs, including motifs evaluated after *in silico* incorporation into the AP restriction-site context. Sequences failing validation are replaced iteratively until all constraints are satisfied.

Optional design extensions include addition of fixed flanking sequences and subdivision of sites into internal positions (`add_positions()`), enabling multi-part barcode architectures within a single restriction-defined site. For position-based designs, *BarcodeEZ* automatically assigns selective 4-mer sticky-end overhangs to internal fragments to preserve ordered restriction cloning compatibility within the Assembly Plasmid. These overhang sequences were selected from a curated set of 4-mers previously benchmarked for high-efficiency directional ligation in restriction-based assembly systems (36) . Structural visualization (`show_structure()`) and export utilities (`write_order_form()`) enable inspection of barcode architecture and generation of synthesis-ready oligonucleotide pool files with appropriate flanking sequences for commercial synthesis.

*LongBarcodeQC: nanopore-based long-read quality control of combinatorial barcode libraries. LongBarcodeQC* (v1.0.0) is a Python (v3.10) command-line package developed for quality control analysis of combinatorial barcode libraries using long-read sequencing data. The software leverages pandas (v2.1.1) for data handling, cutadapt (v5.2) for adapter trimming, parasail (v1.3.4) for semi-global sequence alignment, and seaborn (v0.13.2) and matplotlib (v3.8.4) for visualization within automated HTML reports. *LongBarcodeQC* (v1.0.0 is freely available under the MIT license at https://github.com/ArpiarSaundersLab/LongBarcodeQC and can be installed with pip install LongBarcodeQC ==1.0.0. Documentation: https://arpiarsaunderslab.github.io/LongBarcodeQC/.

*LongBarcodeQC* accepts raw FASTQ reads, a reference plasmid sequence (default: Assembly Plasmid), and a user-defined barcode FASTA file generated by *BarcodeEZ* using the `siteX_posY_bcZ` naming convention. User-configurable parameters include expected insertion number, MCS flanking sequences, restriction enzyme site lists, and barcode-calling thresholds. Base-called reads were adapter-trimmed during primary processing using cutadapt (37) and aligned using minimap2 (38) to both the reference plasmid and the *E. coli* genome, enabling classification of plasmid-derived versus host-derived reads and assessment of plasmid enrichment. Plasmid-aligned reads were converted to a consistent sense orientation based on alignment coordinates. The multiple cloning site (MCS) region was identified within each read by semi-global alignment of predefined 150-bp left and right flanking anchor sequences using parasail, accommodating indel and substitution errors characteristic of nanopore sequencing. The intervening sequence between anchors was extracted as the candidate MCS region for downstream analysis.

Extracted MCS sequences were aligned against the user-provided barcode set using parasail semi-global alignment, and raw alignment scores were computed for each barcode–read pair. Restriction enzyme recognition sequences were detected within each MCS using either a default AP-associated restriction site list or a user-specified set, enabling confirmation of expected restriction site loss or retention during serial cloning and negative selection. Barcode identity was determined by standardizing alignment scores within each read to generate Z-scores across candidate barcodes, and barcode calls were assigned using a user-defined Z-score threshold. *LongBarcodeQC* generates an interactive HTML report summarizing read classification metrics, MCS length distributions, restriction site analyses, and barcode-calling statistics, along with a detailed comma-separated values (.csv) file containing per-read metadata and alignment results. By leveraging full-length nanopore reads and explicit alignment to both plasmid and host genomic references, *LongBarcodeQC* enables molecule-level validation of combinatorial barcode assemblies during iterative AP cloning workflows.

### Reagent sharing

Assembly Plasmids with kanamycin (AP_Kan_) or ampicillin resistance (AP_Amp_) and the described PadlockSeq and TritSeq combinatorial barcode plasmid libraries in either AP_Kan_ or AP_Amp_ will be available from Addgene upon publication (Addgene ID numbers: *Plasmids*: AP-Kan, 261892; AP-Amp, 261893. *Pooled Libraries*: PadlockSeq(AP-Amp), 261888; PadlockSeq(AP-Kan), 261889; TritSeq(AP-Amp), 261890; TritSeq(AP-Kan), 261891). All plasmid sequences are available in Supplementary Data File 1.

### Enzymes

Restriction endonucleases used for combinatorial barcode construction included EcoRI-HF (R3101S), AvrII (R0174S), BamHI-HF (R3136S), KpnI-HF (R3142S), NheI (R3131S), PciI (R0655S), XhoI

(R0146S), SpeI-HF (R3133S), PlutI (R0713S), all from New England Biolabs (NEB), as well as FastDigest MreI (FD2024) and FastDigest MauBI (FD2084) from Thermo Scientific. Exonuclease V (ExoV; M0345S) was obtained from New England Biolabs, and T4 DNA ligase was purchased from Roche (10909246103). All enzymes were used according to the manufacturers’ recommended conditions, except for reaction times detailed below.

### *E. coli* strains and transformation

Bacterial cloning and plasmid propagation were performed using *Escherichia coli* (*E. coli*) strains selected according to experimental requirements. Chemically competent E. coli DH5α cells (Thermo Scientific; EC0111) were used for Assembly Plasmid cloning and routine plasmid preparations. Electrocompetent *E. coli* MegaX DH10B T1R Electrocomp Cells (Thermo Scientific; C640003) were used for all combinatorial barcode cloning in Assembly Plasmids to support high-efficiency transformation of complex libraries. All bacterial strains were cultured and handled according to the manufacturers’ recommended protocols unless otherwise noted.

### Plasmid preparation

Plasmids were prepared using propagation and transformation strategies matched to experimental context. For non-barcoded constructs, chemically competent *E. coli* were transformed by standard heat-shock methods, grown in sterile liquid Luria-Bertani (LB) medium (Invitrogen; 12780-052) supplemented with carbenicillin (Thermo Scientific, J6194906; 100 µg/mL) or kanamycin (Sigma Aldrich, K0254; 100 µg/mL), as appropriate for the plasmid backbone. Non-barcoded plasmid DNA was purified using the ZymoPURE II Plasmid Maxiprep Kit (Zymo Research; D4203) according to the manufacturer’s instructions. For barcoded assembly plasmids, electrocompetent *E. coli* were transformed using a Gene Pulser Xcell Microbial System (1652662; Bio–Rad) with 0.1-cm gap cuvettes (165–2089). Fifty microliters of MegaX cells were transformed with one experimental or control (no barcode insert) ligation reaction and electroporated with the following settings: 2000 V, 200 ohm, 25 uF. Cells were then resuspended in 2 mL of S.O.C medium (15544034, Thermo-Fisher). Recovery suspensions were then transferred into 14 mL culture tubes and incubated at 37 °C for 1 h on a shaking incubator. To assess CFU background rates, a 10 μl sample was removed from experimental and control cultures, serially diluted tenfold in warmed S.O.C medium (ranging from 1:10 to 1:100, 000) and 50 μl samples expanded in duplicate on small-format (10 cm) LB agar plates (see *Estimates of barcode colony forming units* section). To recover complex and uniform barcoded plasmid preparations, 1 mL of each 2 mL cultures from experimental ligations were plated on individual large-format (24 cm × 24 cm) pre-warmed LB agar plates containing the appropriate kanamycin or carbenicillin antibiotic selection. After ∼12 h of growth at 37 °C, bacterial colonies were harvested directly from plates by piecemeal addition of 10-20 mL of LB and gentle dissociation with a cell lifter (Corning CLS3008). Bacteria were then pelleted from the LB media by centrifugation for 10 min at 3, 800 × g at 4 °C, LB supernatant was then removed, and the resulting bacterial pellets were either frozen at -20°C or immediately subjected to plasmid purification. Plasmid purification involved resuspension of bacterial pellets in ZymoPURE P1 (this and all other ZymoPURE buffers adjusted to 28 mL) and processed according to the manufacturer’s guidelines. One column was used per two harvested large-format plates (1-4 g bacterial pellets). Purified DNA was resuspended in Zymo Elution Buffer and quantified using a NanoDrop spectrophotometer (Thermo Scientific). All barcoded plasmid preparations were analyzed through whole plasmid nanopore sequencing for quality control and barcode diversity characterization (see *Nanopore sequencing* and *LongBarcodeQC* sections).

### Molecular methods for serial barcode cloning in the Assembly Plasmid

#### Assembly Plasmid design and synthesis

The Assembly Plasmid (AP) with ampicillin resistance (AP_Amp_) was synthesized *de novo* (Genscript). The design of the AP multiple cloning site was modified from the original STICR design (4 sites)(5), by incorporating two additional sites with distinct restriction-site trios for sticky-end ligation and negative selection (6 sites total). An AP with Kanamycin resistance (AP_Kan_) was cloned in-house via NEBuilder HiFi DNA Assembly Master Mix (NEB Cat #: E2621). PCR amplicons of the AP_Amp_ plasmid (primers, 5’-3’: AP_F1, CTGTCAGACCAAGTTTACTCA; AP_R1, ACTCTTCCTTTTTCAATATTATTGAAG) and the Kanamycin gene (primers, 5’-3’: Kan_F1, AATATTGAAAAAGGAAGAGTATGATTGAACAAGATGGATTGCACGCA; Kan_R1, AGTAAACTTGGTCTGACAGTCAGAAGAACTCGTCAAGAAGG) amplified from pH2B-miRFP670nano (Addgene Accession #: 127438) were generated using Platinum SuperFi II PCR Master Mix (Thermo Scientific Cat #: 12368010) and standard PCR cycling conditions (initial initial denaturation: 98 °C for 30 s; 25 cycles: denaturation, 98 °C for 10 s; annealing, 60 °C for 10 s; extension, 72 °C for 30s; final extension: 72 °C for 5 min).

#### Barcode component ordering and preparation

Single-stranded DNA oligonucleotides encoding individual barcode components were ordered from Integrated DNA Technologies as 5’-phosphorylated oPools using sequences directly generated by *BarcodeEZ* (see *Software Development* section). oPools were resuspended in nuclease-free water (Invitrogen; 10977-015) to final concentrations of 100 µM. oPool component sequences for the TritSeq and PadlockSeq libraries are available in Supplementary Data File 2 and Supplementary Data File 3.

#### AP serial restriction cloning and negative selection

Serial assembly of barcode components into each AP site was performed using restriction enzyme cloning coupled to enzymatic negative selection to deplete plasmids with failed barcode ligation events. Single-stranded barcode oPools were annealed to generate double-stranded fragments with 5’ overhangs by combining equimolar forward and reverse oPools (1 µL each; 100 µM) with 5 µL 10× T4 DNA ligase buffer and nuclease-free water to a final volume of 50 µL, followed by incubation at 94 °C for 2 min and gradual cooling to room temperature. Annealed barcode fragments were aliquoted and stored at −20 °C prior to use. For each barcode insertion site, 1 µg AP backbone was digested in a 50 µL reaction using the site-appropriate restriction enzyme pairs in 1× rCutSmart buffer (NEB) for 8 h at 37 °C. Digested backbones were resolved by agarose gel electrophoresis. The linearized AP plasmid band was excised and purified prior to ligation. Ligation reactions (final volume 50 µL) were assembled with 500 ng of digested AP backbone (final concentration of ∼7.8 nM), 1 µL of annealed barcode fragments (diluted to 0.05 -1 nM), 1 µL T4 DNA ligase, and 1× ligation buffer and incubated at 16 °C for 2 h. Vector-to-insert molar ratios ranged from 1:0.3 to 1:5 during optimization, and were set at 1:0.3 and 1:5 for the assembly of the PadlockSeq and TritSeq libraries, respectively. Control ligations lacking annealed barcode fragments were generated in parallel for transformation to calculate background colony forming unit (CFU) rates (see *Estimates of barcode colony forming units* section). Ligation and control ligation products were purified using silica column–based DNA cleanup (Zymo DNA Clean & Concentrator-5; Zymo Research, D4003), eluted in 10 µL of Elution Buffer, and subject to a secondary restriction digest targeting a negative-selection site flanked by the restriction sites used for barcode ligation, using 10 U of restriction enzyme in 1× rCutSmart buffer (50 µL total volume) for 8 h at 37 °C to selectively cleave the unmodified AP backbone site. Negative selection reactions were column purified and eluted in 10 µL of Elution Buffer before transformation into electrocompetent *E. coli* and subsequent plasmid preparation (see *Plasmid Preparation* section). Site-specific plasmid preparations were evaluated with nanopore sequencing and *LongBarcodeQC* analysis (see *Nanopore sequencing* and *LongBarcodeQC* sections). Plasmid preparations that passed sequencing-based quality control metrics were used as input for the next site in the restriction cloning and negative-selection workflow, enabling site-wise serial assembly of high-complexity combinatorial barcode plasmid libraries.

#### Estimates of barcode colony-forming units

For each site-wise barcode cloning step in the AP and for the expression vector transfer cloning, estimates of the total number of *E. coli* transformants with barcoded plasmid were made based on serial dilutions (1:10-1:100, 000) of transformation products from ligations with and without (control) barcode fragments (described in *Plasmid preparation* section). Colony-forming units (CFUs) were counted by hand across dilution conditions. To estimate the number of barcoded plasmids harvested in total from collections of large agar plates, background CFUs were subtracted from the total CFUs estimated from the transformation volumes spread on the large plates. This method assumes a conservative one-to-one correspondence between a transformant and plasmid and thus likely represents a lower bound on barcode diversity in each plasmid library (39) .

#### Agarose gel electrophoresis and DNA extraction

Agarose gel electrophoresis was used to monitor AP plasmid size and integrity during serial assembly, as well as to isolate linearized DNA for downstream cloning. AP plasmid size and integrity were monitored on 1% agarose gels (VWR, 97062-244) prepared in 1x TAE buffer and run at 100 V for 30-60 min and visualized using SYBR Safe DNA stain (Thermo Fisher, S33102) under blue-light illumination. DNA for downstream cloning was resolved on 1% low-melt agarose gels (Sigma-Aldrich, A9414) prepared in 1x TAE buffer and run at 65 V for 1-2 h, followed by excision of the corresponding gel bands. For transfer of fully assembled barcodes, a restriction fragment containing barcode MCS was similarly excised following restriction digest of a pair of AP transfer sites (see *Transfer Cloning* section). Excised gel slices were processed for DNA recovery using the Zymoclean Gel DNA Recovery Kit (Zymo, D4001) according to the manufacturer’s instructions, and DNA eluted in 10 μL of Elution Buffer was quantified using NanoDrop spectrophotometer before downstream cloning steps.

### Molecular methods for barcode transfer cloning into a recipient expression vector

*Generation of the N2c Rabies Virus Genome Plasmid with a Transfer Site* CVS-N2c-nl.EGFP-SypGFP was acquired from Addgene (Accession #: 172380). Primers targeting the 3’ UTR region of nl.EGFP were used to introduce the Transfer A (MreI) and Transfer B (MauBI) sites flanking the negative selection site (TspMI) and ordered as ultramers from IDT (5’-3’: CVS_c18_v2_TspMI_TermBv2_F1, ACCTCCCGGGACCGGCGCCGGCGGCTAGACATGAAAAAAACTAACACTCCTCCGGTACCGCC A and CVS_c18_v2_TspMI_TermAv2_R1, ACCTCCCGGGACTGCCGCGCGCGGCCCCCTCGACGAATTCCAAATGTGGTATGGCTGATTATG). The PCR product was gel extracted and subjected to a procedure for template DNA removal (DpnI digest), TspMI overhang generation and ligation using the following protocol: In a 20 μL final volume, PCR product DNA (up to 500 ng) was combined with 2 units of DpnI (1 ul), 2 μL of 10x rCutSmart Buffer (NEB) and incubated for 1 hr at 37 °C. The resulting reaction was column cleaned, eluted in 10 μL of Elution Buffer, and added to a 50 μL reaction containing 10 units of TspMI (2 ul) and 5 μL of 10x rCutSmart Buffer (NEB) and incubated at 75 °C for 1 h. The reaction was column cleaned, eluted in 10 μL of Elution Buffer and DNA quantified using a spectrophotometer. 20-500 ng of DNA was used as input to a ligation reaction in a total of 30 ul, containing 1 unit T4 DNA ligase (1 μL), 3 μL of 10x ligase buffer (Roche) and incubated at 4 °C for 2 h. 2 μL of ligation reaction was used for transformation. The DpnI and restriction digest reactions can be combined if compatible. Of note, the PadlockSeq combinatorial barcodes were transferred in the reverse-complement orientation as compared to the AP, such that each site’s conserved sequence was more proximal to the polyadenylation signal. The resulting plasmid was called “CVS-N2c(deltaG)-N2c-nl.EGFP-SypGFP-TermBAv2_TspMI” (Supplementary Data File 1).

#### Transfer cloning

Transfer cloning of the PadlockSeq or TritSeq MCS from the AP to a second AP (with orthogonal antibiotic resistance) or Transfer-site modified expression vector began with a restriction digest to free the barcode-containing MCS. In a 20 μL reaction, 1 μg of AP with completed barcodes was combined with 1 μL of FastDigest MreI, 1 μl of FastDigest MauBI, 2 μl of 10x FastDigest Green Buffer and incubated at 37 °C for 8 h and then subjected to agarose gel electrophoresis and gel extraction. Transfer plasmid (1 μg) was subjected to an equivalent restriction digest and gel extraction. Ligation was performed in a 50 µL reaction containing 500 ng of digested transfer plasmid DNA and barcoded MCS DNA at a 1:3 vector:insert molar ratio. Ligation reactions were incubated at 16 °C for 4 h, followed by column cleaning (1 column per reaction) and eluted in 10 μL elution buffer. To remove any transfer plasmids with an intact transfer site, cleaned ligation product was digested in a 50 μL reaction containing, 10 units of SmaI (0.5 μL) (same recognition site as TspMI) and 5 μL of 10x rCutSmart Buffer (NEB) incubated at 37 °C for 8 h. Digest reactions were subsequently column cleaned, eluted in 10 μL of elution buffer and transformed.

#### N2c rabies virus plasmids with single PadlockSeq barcodes

To select example CVS-N2c-nl.EGFP-SypGFP_PadlockSeq_ transformants with single PadlockSeq barcodes, 20 individual clones were sampled from large agar plates, maxiprepped and sequence confirmed. Six plasmid preparations harboring unique PadlockSeq barcodes A-F were selected for rabies virus packaging (Supplementary Data File 1).

### Packaging of G-deleted N2c Rabies Virus

Packaging of six distinct CVS-N2c-nl.EGFP-SypGFP_PadlockSeq_ barcoded genomes (lettered A-F) with native N2c glycoprotein followed the protocol established by Sumser et al (40). HEK293-GT cells were used to rescue CVS-N2c rabies viral vectors. Cells were seeded in 6-well culture plates and grown to approximately 80–90% confluence prior to transfection. Cells were then transfected with the rabies vector plasmid together with the N2c helper plasmids (pCAG-N, pCAG-P, pCAG-L) and pCAG-t7pol using xfect transfection reagent in optiMEM media. OptiMEM media was replaced 6 h post-transfection with DMEM supplemented with 10% FBS. Twenty-four hours after transfection, cells were resuspended and re-plated into 100 mm culture dishes and incubated at 37 °C with 5% CO₂ until full confluence was re-established. Cultures were maintained with frequent media changes until approximately 100% of cells exhibited fluorescence, typically 5–6 days post-transfection and 1–2 days after fluorescence was first observed. At this stage, culture supernatant was collected, filtered, concentrated, aliquoted, and stored at −80 °C. N2c rabies virus titers were assessed using HEK cells following the protocol from Wickersham et al (41). One day after plating, cells from one well were resuspended and counted, while cells in the remaining wells were transduced with 1 µL of concentrated virus in serial dilutions (using 1x PBS) ranging from 1:10 to 1:10, 000. Three days post-transduction, cells were resuspended, washed with 1x PBS, and fixed with 4% paraformaldehyde (PFA) for 10-15 min at room temperature. The fraction of GFP+ transduced cells was quantified using flow cytometry (BD FACSymphony Analyzer A5). Thresholds for positive events were defined using untreated control cells. Viral titers were calculated using the following formula: Titer (IU/mL) = −ln(P_uninfected) × n/v, where P_uninfected is the fraction of uninfected cells, n is the number of total cells in each well at the time of viral transduction, and v is the volume in milliliters of the viral stock added. Wells with between 1-10% of fluorescent cells were used to calculate titers, based on an average of n=2 technical replicates.

### Transduction of PadlockSeq combinatorial barcodes into the mouse brain

#### Animal husbandry

All mouse procedures were approved by the Oregon Health & Science University (OHSU) Institutional Animal Care and Use Committee (protocol #IP00003776) and conducted in accordance with institutional and federal animal care guidelines. Experimental animals were C57BL/6 mice derived from breeding crosses maintained at OHSU.

#### Stereotactic injection

Stereotactic injections were performed on adult C57BL/6 mice (P50–P60) using standard aseptic surgical procedures. Animals were anesthetized with isoflurane (induced at 3–4% and maintained at 1–2%) and secured in a stereotactic frame; body temperature was maintained throughout surgery. Bilateral craniotomies were made over the primary somatosensory cortex (S1), targeted using stereotactic coordinates relative to bregma (anteroposterior: approximately −1.0 mm; mediolateral: approximately ±2.5 mm; dorsoventral: approximately −0.5 to −0.7 mm from the cortical surface). CVS-N2c-nl.EGFP-SypGFPPadlockSeq(G) viruses (variants A–F) were mixed at equal titers immediately prior to injection, and a total volume of 500 nL per hemisphere at a final concentration of 1 × 10⁸ IU/mL was delivered using a pulled glass micropipette at an infusion rate of 50 nL/min. Following injection, the pipette was left in place briefly before slow withdrawal to minimize reflux. The scalp was sutured, animals received ketoprofen for postoperative analgesia, and mice were monitored daily for at least 5 days following surgery to assess recovery and overall health.

### *In situ* sequencing of PadlockSeq barcodes

#### Brain section preparation

All tissue processing was performed under biosafety level 2 (BSL-2) conditions. Animals were transcardially perfused with ice-cold dissociation buffer (in mM: 82 Na₂SO₄, 30 K₂SO₄, 10 HEPES, 10 glucose, and 5 MgCl₂), after which brains were rapidly dissected and flash-frozen on aluminum foil over a liquid nitrogen–ethanol slurry. Frozen tissue was embedded in optimal cutting temperature (OCT) compound using standard molds and stored at −80 °C until sectioning. Brain sections were prepared at 50 µm thickness using a cryostat (Leica Microsystems, CM3050) under standard cryosectioning conditions. Sections were examined for GFP fluorescence in and around primary somatosensory cortex to identify individual sections with dense rabies virus labeling. Selected sections were transferred to a 12 well glass bottom plate and processed for *in situ* sequencing using a precommercial version of the Pyxa platform following manufacturer’s guidelines (Stellaromics).

#### In situ sequencing on the Pyxa platform

Spatially resolved transcriptomic profiling was performed using the STARmap workflow (18). Sections were pretreated and permeabilized before hybridization with SNAIL Probes in Hybridization Buffer using a custom 18-probe panel targeting sites 1-3 of the PadlockSeq A-F barcodes (Supplementary Data File 4). Unbound probes were removed with Post-Hybridization Wash Buffer. Hybridized probes were subsequently ligated and amplified *in situ*, then embedded in hydrogel to preserve spatial architecture. Remaining tissue was cleared with Clearing Buffer and Clearing Enzyme, and sample quality was verified using the Sample Verification Solution. Multiple sequencing rounds were imaged with DAPI to label cell nuclei and fluorescent probes to detect amplicons representing individual transcripts. Sequencing rounds were imaged on a high-resolution confocal microscope with z-stack acquisition and processed using a precommercial Pyxa^®^ instrument to capture three-dimensional distributions of PadlockSeq RNA barcode decodes.

#### Pyxa data analysis

PadlockSeq barcode A-F analysis was performed using custom analysis scripts written in Python using a Pyxa output file consisting of x, y, z decode positions as input. Analysis consisted of three major steps: decode filtering, cell assignment, and cell-resolved expression analysis. Probe-specific decode filtering was performed downstream of a custom permutation-based sampling approach to score spatially clustered versus non-clustered decodes. Spatially clustered decodes were retained and assigned into single cells through spatially resolved Leiden-based clustering. The subsequent cell assignments were then used to assess site-specific decode levels across sites 1-3 of the PadlockSeq combinatorial barcodes. All analysis scripts are available at https://github.com/ArpiarSaundersLab/2026_combinatorial_barcode_analysis.

### Nanopore sequencing

Nanopore sequencing was performed either in-house or through third-party commercial services (GeneWiz or Plasmidsaurus) using Oxford Nanopore Technologies (ONT) platforms. For in-house sequencing, plasmid DNA was purified using Zymo DNA Clean & Concentrator-5 columns and DNA concentrations were assessed using a NanoDrop spectrophotometer (Thermo Scientific) prior to library preparation. Sequencing libraries were prepared using the Rapid Sequencing Kit v14 (SQK-RAD114; ONT) according to the manufacturer’s instructions using the recommended DNA input (50 ng), followed by transposase-mediated tagmentation and rapid sequencing adapter attachment. Sequencing was performed on Flongle flow cells (R10.4.1; FLO-FLG114) using the Flongle Sequencing Expansion Kit (EXP-FSE001) on a MinION Mk1B device, with flow cells primed and libraries loaded according to ONT protocols. Sequencing runs were initiated and monitored using MinKNOW software (v6.10.12) with default settings optimized for Flongle flow cells, and base-calling was performed using Dorado software (v7.13.6; ONT). Reads were filtered using standard quality thresholds and FASTQ files were used as input into *LongBarcodeQC* (see *Software Development* section).

### Data analysis, plotting and statistical testing

#### General

Figure plots were generated from *LongBarcodeQC* output files using the R programming language. Key analysis packages included dplyr(1.1.4), tidyverse(2.0.0), ggplot2(4.0.1), and ggsankey(0.0.99999). Statistical analyses were performed in R. All analysis scripts are available at https://github.com/ArpiarSaundersLab/2026_combinatorial_barcode_analysis.

#### Combinatorial barcode diversity

Barcode diversity was quantified from whole-plasmid long-read sequencing data using a pore-aware approach to mitigate duplex sequencing artifacts. For each barcode site/position, barcode collisions were quantified using only pairs of reads originating from different nanopore channels, thereby excluding read pairs that could represent duplicate sequencing of the same DNA molecule. The observed collision frequency among different-pore read pairs was then adjusted for the fraction of all possible read pairs arising from different-pore comparisons, calculated from the empirical distribution of reads across nanopore channels. This yielded the pore-corrected collision probability *λ*^^^ , representing the probability that two randomly sampled molecules contain the same barcode element. Inverse Simpson diversity was calculated as 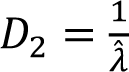 Simpson evenness was defined as 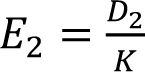, where *K* is the number of possible barcode elements at that site/position. Independent assortment between barcode sites was assessed using pairwise mutual information (MI). Finite-sampling bias was estimated from 1, 000 random permutations of barcode identities between sites, and permutation-adjusted normalized MI (NMI) was calculated as 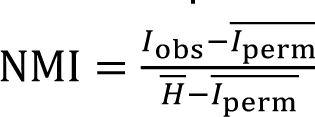, where *H* is the mean entropy of the two sites. Full combinatorial barcode diversity was assessed using the same collision-probability estimator, treating each unique combination of site/position barcode identities as a single barcode. The corresponding inverse Simpson diversity 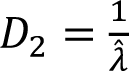 represents the effective number of equally abundant combinatorial barcodes. The 95% confidence intervals related to barcode abundance percentages in Figure 8D were calculated by multinomial sampling of a uniform library at the observed depth: 2, 000 replicates were drawn as (*n*_1_, … , *n_v_*) ∼ Multinomial(*N*, 1/*v*, … , 1/*v*), barcode abundances were sorted in decreasing order within each replicate, and the 2.5th and 97.5th percentiles across replicates were calculated at each rank.

## RESULTS

### Overview of the workflow for combinatorial barcode design, assembly and quality control

To assemble and quality control plasmid libraries encoding diverse and flexible combinatorial barcode designs, we developed a four-step workflow using purpose-built computational tools and DNA cloning plasmids integrated with long-read nanopore sequencing analysis (**Figure 1**). Three components facilitate assembly and quality-control of user-designed combinatorial barcode libraries. A fourth component facilitates the transfer of assembled combinatorial barcodes into expression vectors for experimentation. Here we summarize this workflow and provide a roadmap for how each component is reported in the results.

The first two components consist of 1) purpose-built “Assembly Plasmids” (APs) for optimized for serial restriction cloning and expression vector transfer and 2) a pip-installable Python package called *BarcodeEZ* developed for barcode design. Since the structure of the AP is critical for serial barcode design and assembly, the logic of AP design is introduced first (**Figure 2**). Next, we introduce how *BarcodeEZ* design software uses object-oriented functionality and the AP sequence structure to design combinatorial barcodes for error-free assembly (**Figure 3**).

**Figure 2.**
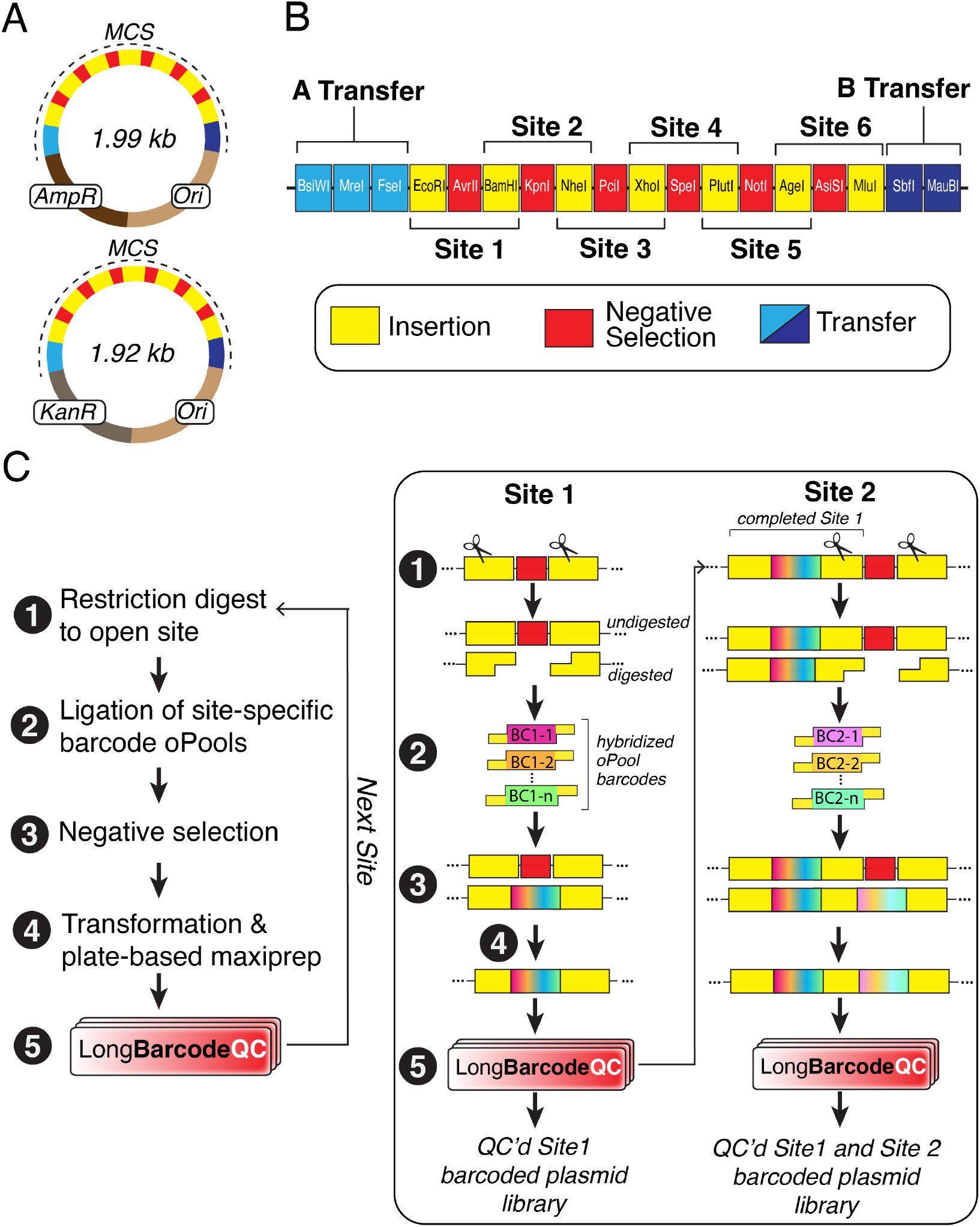
Assembly Plasmid design supports high-fidelity serial barcode cloning. **A.** Schematic maps of Assembly Plasmids (APs) with either ampicillin (AP_Amp_) or kanamycin (AP_Kan_) antibiotic resistance. Each AP includes a custom multiple cloning site (MCS) arranged as alternating insertion (yellow) or negative-selection (red) restriction sites, flanked by additional restriction sites for transfer cloning of the fully assembled combinatorial barcode (light and dark blue). **B.** Schematic of the detailed layout of the MCS. Six sites containing restriction site “trios” are available for assembly. Each trio consists of a pair of 6-bp restriction enzyme recognition sequences for barcode ligation (“Insertion”), flanking an additional restriction sequence used to digest and remove plasmids with insertion failures (“Negative Selection”). Adjacent sites share insertion sequences. For example, the right insertion sequence for site 1 (BamHI) is the left insertion sequence for Site 2. After assembly, 8-bp restriction sequences flanking sites 1-6 are used to transfer the completed combinatorial barcode to a desired EV through restriction cloning (“Transfer A” and “Transfer B”). **C.** Workflow for site-wise serial cloning in the AP. Each numbered step is generally described at left and schematized in more detail at right using site 1 and 2 as examples.

**Figure 3.**
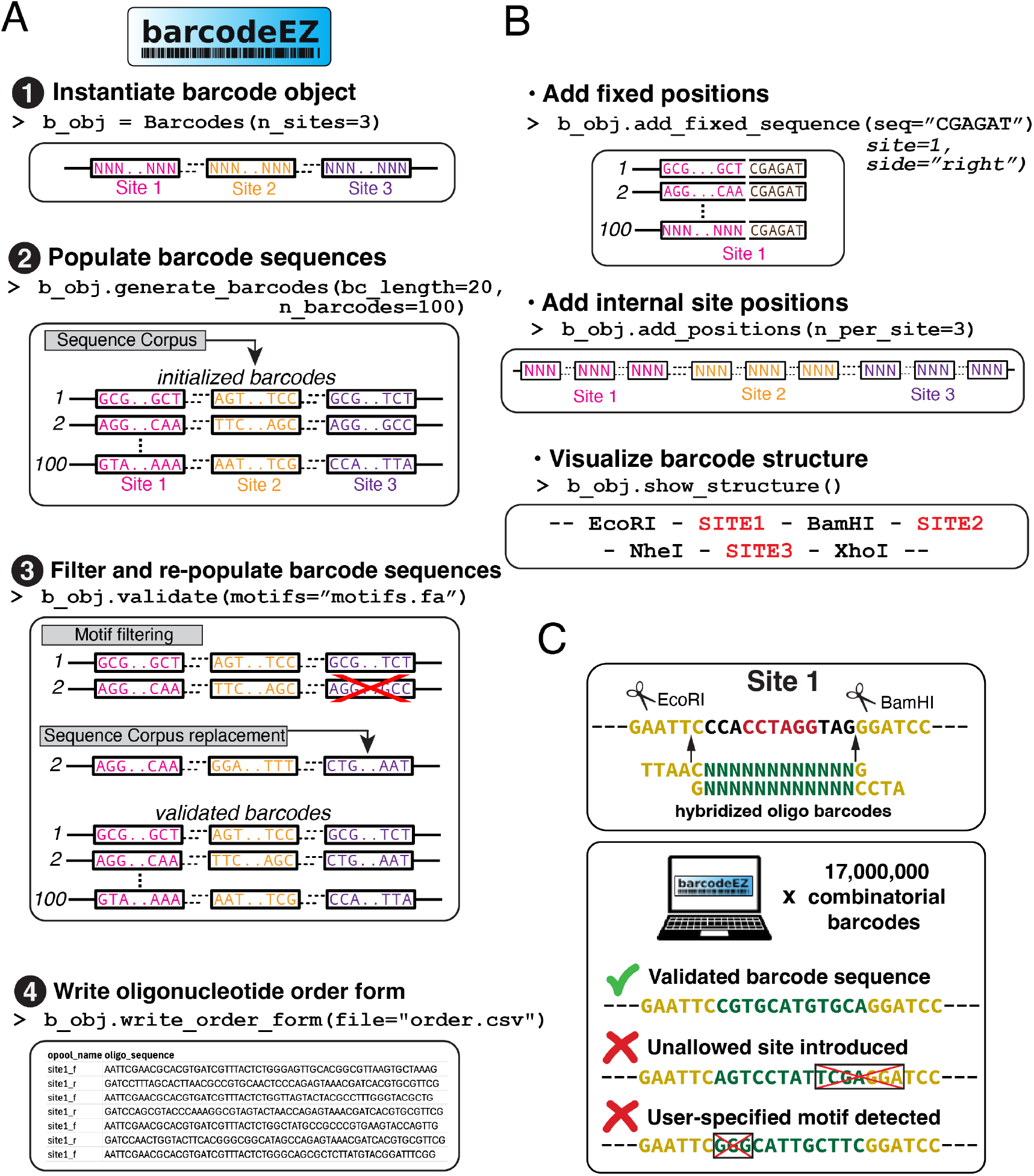
*BarcodeEZ*: a software package for combinatorial barcode design and site-aware assembly. **A.** Ordered workflow of the four minimal *BarcodeEZ* methods to generate a user-specified combinatorial barcode library to minimize assembly errors. For each method, the purpose, example call, and a schematic representation are shown. **B.** Examples of optional *BarcodeEZ* methods to further customize combinatorial barcode designs. Such methods enable addition of fixed sequences, subdivision into site-internal ‘positions’ to expand combinatorial diversity, and visualization of designed barcode structures. **C.** Detailed schematic of the `.validate()` method process. *BarcodeEZ* models the insertion of each barcode element into the desired site. Events that introduce allowed sequences are retained, while events that create disallowed restriction sites or unwanted user-defined sequence motifs are flagged and replaced with updated barcode sequences from the corpus that pass user-defined filters.

To demonstrate generalizability of the AP/*BarcodeEZ* components, we next present two unique combinatorial barcode architectures purpose-built for distinct types of probe-based decoding (“TritSeq” and “PadlockSeq”; **Figure 4**). To illustrate the validation process for new combinatorial barcode designs before full assembly, we deliver example PadlockSeq barcodes to the mouse brain with a viral vector and use probe-based *in situ* RNA sequencing to demonstrate robust decoding of PadlockSeq barcode identities (**Figure 5**).

**Figure 4.**
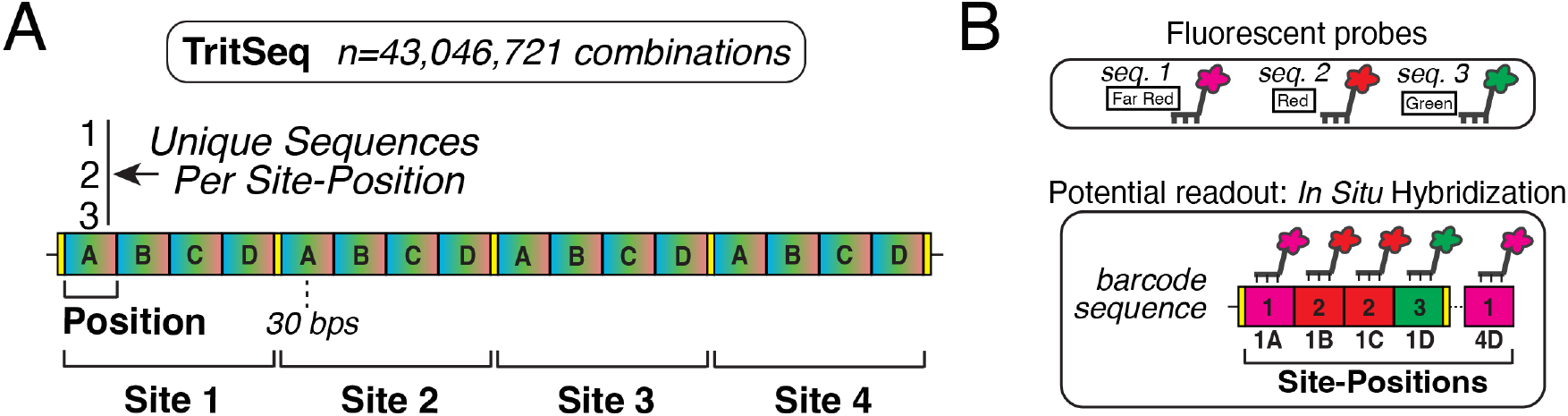
TritSeq combinatorial barcode architecture. **A.** Schematic overview of the TritSeq combinatorial barcode. The TritSeq design illustrates a design amenable to single-probe, three-channel decoding. The TritSeq barcode consists of n=16 30-bp variable sequences, organized as four adjacent positions (A-D) within each of four assembly sites (1–4). The TritSeq barcode comprises 16 30-bp variable sequences organized as four adjacent positions (A–D) within each of four assembly sites (1–4), encoding 3^16^ =43, 046, 721 unique sequences that can be read out with 48 probes. The complete barcode is 552 bp long (complete MCS = 631 bp). **B.** Example probe-based decoding strategy based on three spectrally distinct fluorescent *in situ* hybridization probes targeting each TritSeq site/position.

**Figure 5.**
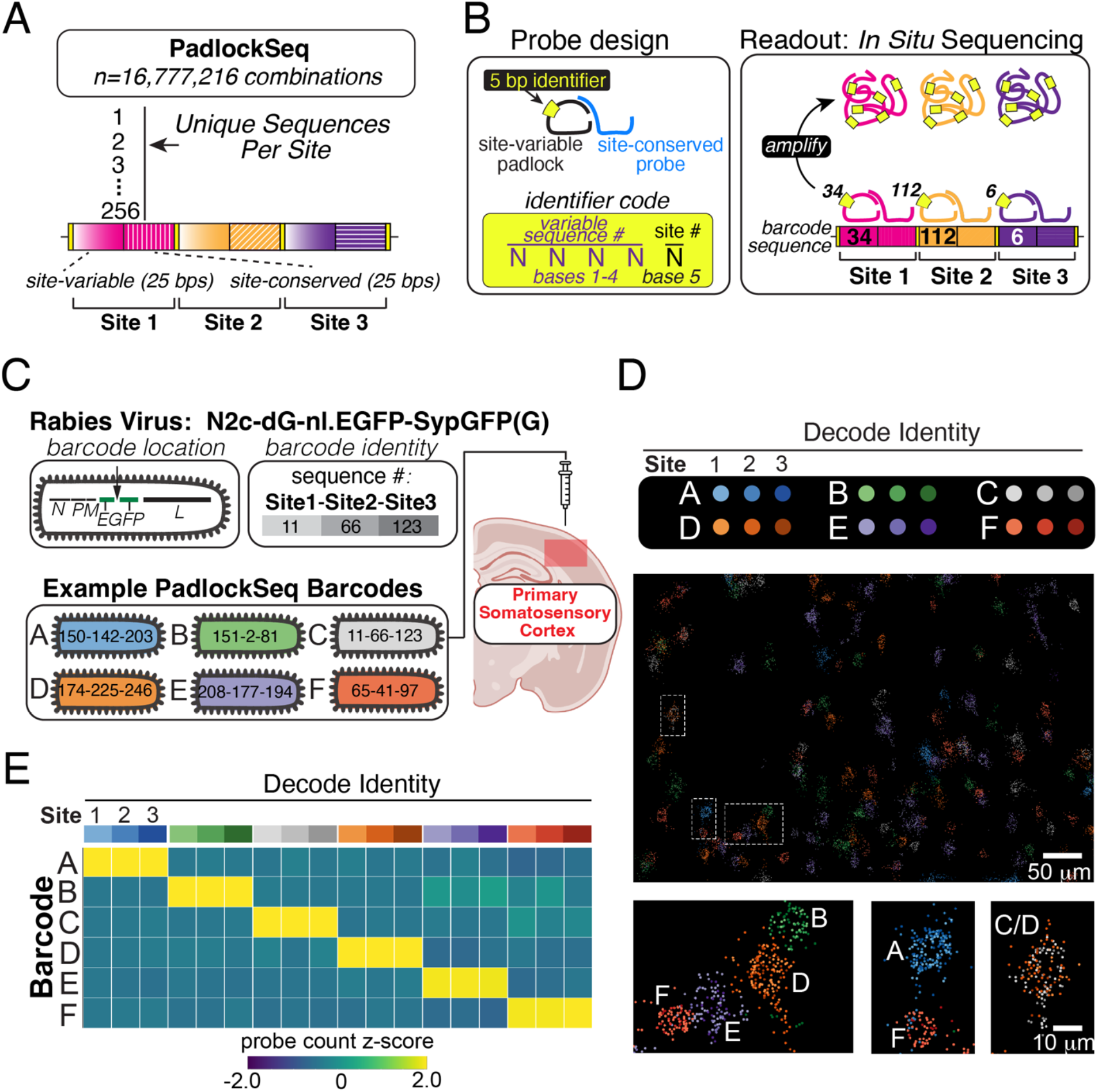
Design and *in vivo* validation of the PadlockSeq combinatorial barcode architecture using probe-based *in situ* sequencing. **A.** Schematic overview of the PadlockSeq combinatorial barcode, designed for two-part padlock probe decoding. The PadlockSeq barcode consists of three adjacent sites. At each site, one of 256 site-variable 25-bp sequences is positioned immediately adjacent to a site-conserved 25-bp region. The PadlockSeq barcode design encodes 256^3^ = 16, 777, 216 unique sequences designed for readout with 768 probe pairs and is 162 bp in length (complete MCS = 276 bp). **B.** Left, probe-based decoding strategy using STARmap. PadlockSeq mRNA barcodes are detected with a two-part probe pair. Site-specific primer probes each bind to their respective site-conserved sequence, while a library of n=768 padlock probes bind to each site-variable sequence. Each padlock probe has a unique 5 bp identifier that is sequenced *in situ* using the “sequencing with error-reduction by dynamic annealing and ligation” (SEDAL) method. Bases 1-4 of the probe identifier indicate the number of the site-variable sequence (barcodes 1-256), while base 5 indicates the specific site (sites 1-3). Right, rolling circle amplification events amplify probe identifier sequences for SEDAL. **C-D.** *In vivo* detection of PadlockSeq mRNA combinatorial barcodes in mouse brain after delivery with a glycoprotein-deleted N2c strain rabies virus expression vector followed by probe-based *in situ* SEDAL sequencing using the Pyxa platform. **C.** Experimental schematic. Six PadlockSeq barcodes (lettered A-F), each containing unique site-variable sequences at all three sites, were independently introduced into the rabies virus N2c-dG-nl.EGFP-SypGFP genome at the indicated location (**Methods**). Each rabies virus variant was packaged independently into virions with the native N2c glycoprotein, enabling direct infection of neurons without additional cellular transmission. Virions A-F are colored-coded and the variable sequences present at each site are listed and separated by dashes. Virion A-F were mixed at equal titers and injected intracerebrally into the primary somatosensory cortex, directly transducing cortical neurons. The red box on the coronal brain schematic indicates the imaging region in primary somatosensory cortex. **D.** Z-projection through a 50 μm brain section showing the location and identities of *in situ* sequencing events (“decodes”) corresponding to probe-initiated rolling circle amplification from 264 rabies-infected neurons (**Methods**). Decode colors indicate each A-F variable sequence barcode number and the shade reflects the site of origin. The three insets below show seven example neurons highlighting robust detection of site 1-3 decodes in each single cell. Six neurons express single PadlockSeq barcodes (A, B, D, E, and F), while a seventh expresses two PadlockSeq barcodes (C and D). **E.** Heat map of site-specific decode counts across the 264 segmented neurons (**Methods**). Site-specific probe counts (x axis) are organized by expected relationships in PadlockSeq barcodes A-F (y axis). Site-specific decode values represent a Z-score based on counts across all segmented cells.

Serial assembly of full combinatorial barcode libraries benefits from immediate and quantitative quality control to avoid the accumulation of barcode cloning errors. To fill this need and evaluate stepwise cloning outcomes in a fast and cost-effective manner, we introduce the third component, a pip-installable Python package called *LongBarcodeQC. LongBarcodeQC* is custom-built software for analysis of whole plasmid nanopore sequencing data in an AP-and barcode sequence-aware fashion, generating standardized outputs for user-friendly evaluation of cloning outcomes. After introducing core *LongBarcodeQC* functionality (**Figure 6**), we use *LongBarcodeQC* analysis to validate how features of the AP design enable expected barcode cloning outcomes with high efficiency (**Figure 7**). With assembly and quality-control components introduced, we summarize the stepwise assembly and quality-control of the TritSeq and PadlockSeq combinatorial barcode libraries in the AP (**Figure 8**). Finally, we illustrate the fourth component – the transfer of completed combinatorial barcode sequences from the AP to an expression vector (EV) of interest. We do this by detailing how the EV is modified in a fast and flexible way, followed by a molecular protocol to transfer combinatorial barcodes and assess barcoded EV with *LongBarcodeQC*, using introduction of PadlockSeq combinatorial barcodes into a glycoprotein-deleted rabies virus genome plasmid as an example (**Figure 9**).

**Figure 6.**
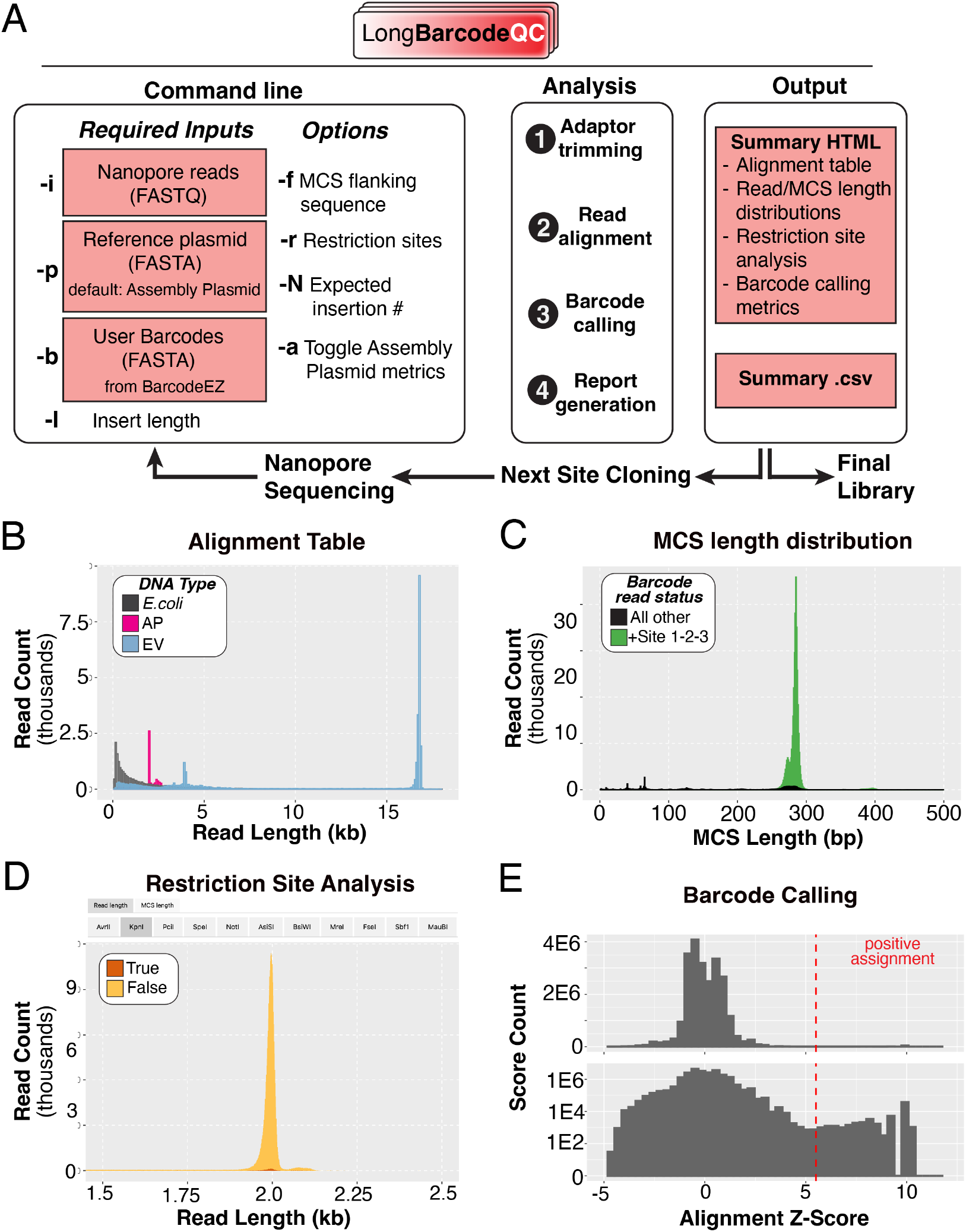
*LongBarcodeQC*: A software package for combinatorial barcode analysis using whole plasmid long-read DNA sequencing. **A.** Overview of *LongBarcodeQC* functionality and workflow. Left, command-line options for required inputs (colored boxes) and optional arguments. Center, ordered analysis steps: Following adapter trimming, reads are aligned to the reference plasmid and marked as plasmid, bacterial, or of unknown origin. Barcode calling is performed by aligning expected barcode sequences to the MCS region of plasmid reads and assigned using a Z-score threshold based on these alignment scores (set by default or provided by user). Right, *LongBarcodeQC* output files. The first output file is an interactive report in HTML format that includes alignment results, plots of read and MCS lengths, barcode calling metrics and restriction site detection. The second output is a CSV summary file that includes verbose metadata associated with each analyzed read, supporting additional downstream analysis. **B-E.** Example descriptive plots from the HTML summary file. **B.** Histogram of analyzed read lengths, color-coded by origin (AP plasmid, magenta; Expression Vector, blue; *E. coli* genome, dark grey). **C.** Histogram of length distributions for computationally-extracted MCS regions from each read, color-coded by the number of barcodes called at each site. **D.** Interactive restriction site analysis. Read length histogram color-coded by exact matches for restriction sites provided by the user (-r). **E.** Histogram of the barcode alignment scores for each read, shown with a linear (top) and log-scaled y axis (bottom). The Z-score threshold for barcode assignment is shown (red dotted line).

**Figure 7.**
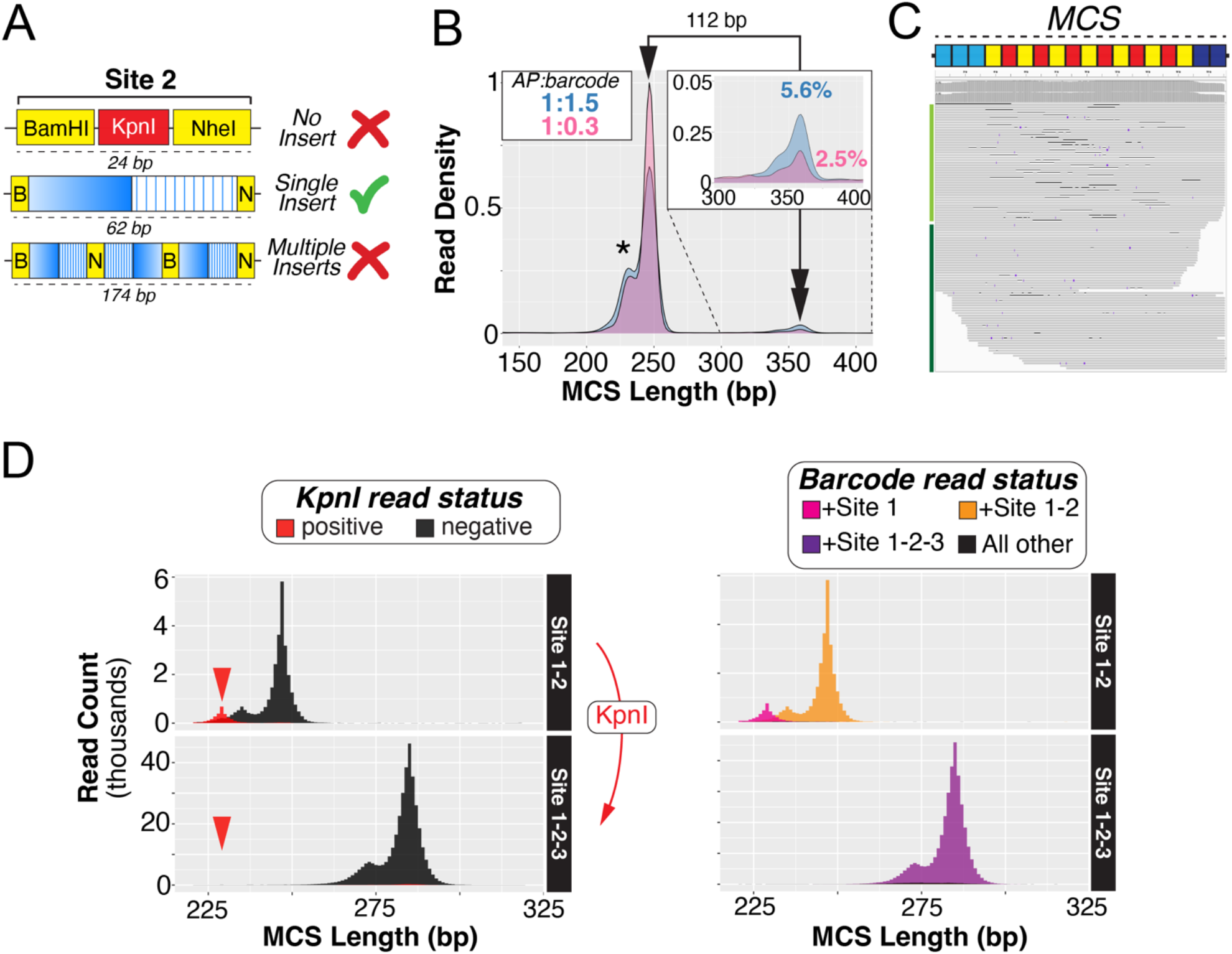
Quality control of serial barcode cloning in the Assembly Plasmid as assessed by LongBarcodeQC. **A.** Schematic of site-specific barcode cloning outcomes using PadlockSeq site 2 as an example. Each site-specific cloning event results in either a failed barcode ligation (“No Insert”), a single barcode ligation (“Single Insert”), or multiple barcode ligations (“Multiple Inserts”). *LongBarcodeQC* uses the length and sequence composition of plasmid long-reads to discriminate between desired “Single Insert” and undesired “No Insert” or “Multiple Insert” events. **B.** Comparison of “Multiple Insert” events for two different site 2 PadlockSeq barcode ligation conditions. Density plot of MCS lengths color-coded by ligation condition (AP:barcode fragment molar ratios 1:1.5, blue, n = 36, 743 reads; 1:0.3, purple, n = 39, 146 reads). The expected MCS length for single-insert events is 247 bp (arrowhead). The double arrowhead and inset highlight a second MCS length peak at 359 bp, corresponding to multiple barcode insertions. The observed 112-bp separation between MCS length peaks (359 − 247 bp) matches the expected 112-bp length difference between Single and Multiple Insertion products (174 − 62 bp). The shorter mode in the MCS length distribution (asterisk) reflects nanopore sequencing deletions (Panel C). Different vector:insert molar ratios affect the number of Single versus Multiple barcode insertion events (Chi-square test, p < 1e-16; odds ratio = 2.3, Fisher’s exact test). **C.** Occasional nanopore sequencing deletions distort inferred MCS lengths. Example reads from the population in the shorter mode (asterisk in panel B) aligned to the AP MCS. Reads with internal (light green bar at left) or end-terminal (dark green bar at left) deletions (shown with thinner black lines) reduce estimated MCS lengths (15 bp on average in this example). **D.** Validating the efficacy of negative selection of No Insert plasmids using PadlockSeq site 2 to site 3 cloning as an example. Histogram of MCS lengths after site 2 barcode cloning (top, n = 35, 771 reads), followed by KpnI digest (negative selection for site 2) and subsequent site 3 barcode cloning (bottom, n = 326, 324 reads). Left, reads are color-coded by the presence of the KpnI site. Right, reads are color-coded by sites 1, 2 and 3 barcode calls. Negative selection with KpnI removes APs lacking a site 2 barcode insertion (Chi-square test, p < 2.2e-16; odds ratio = 0.003, Fisher’s exact test). Red arrowheads indicate the expected MCS length for plasmids that lack a site 2 barcode insertion (18 bp shorter).

**Figure 8.**
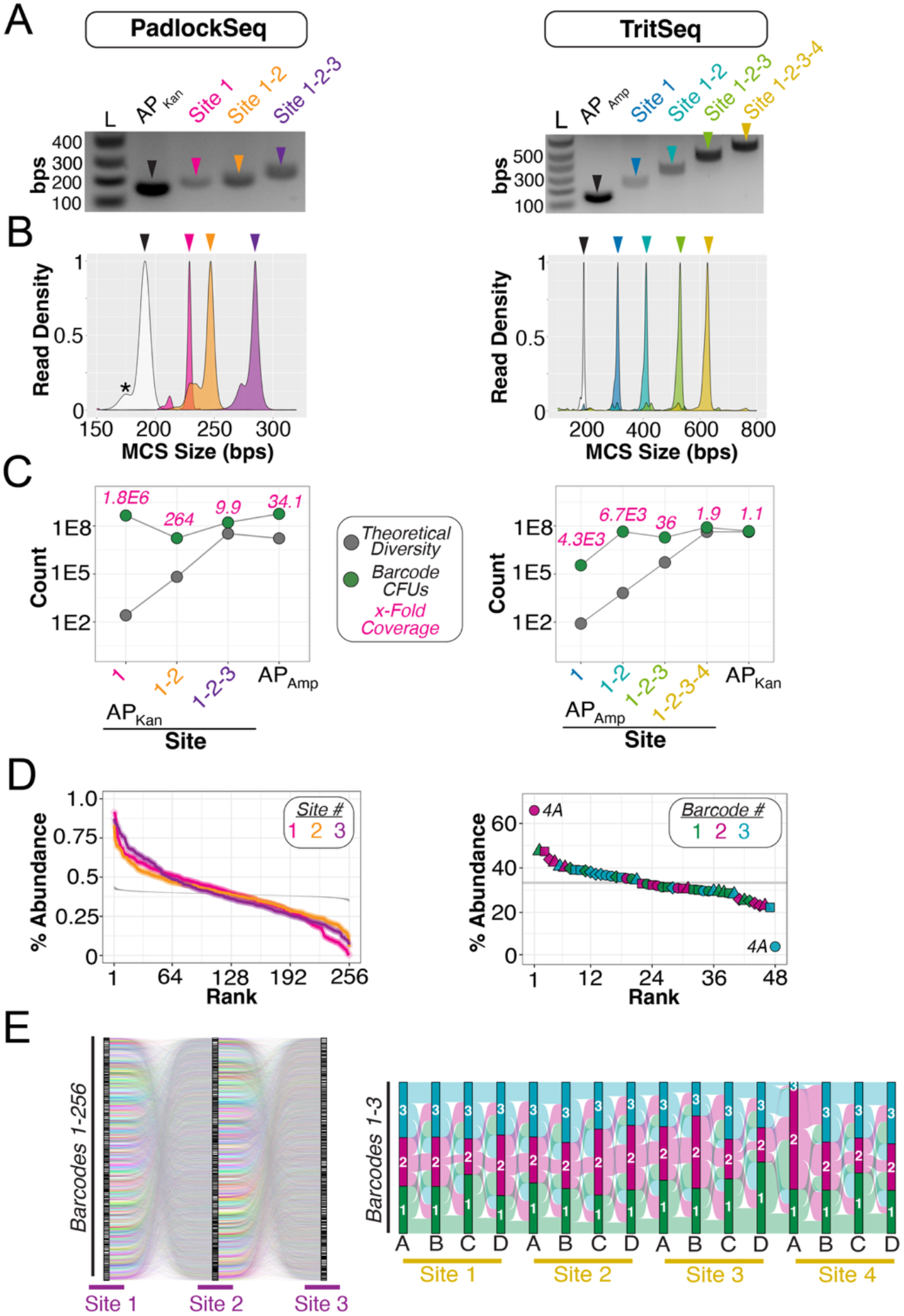
Serial assembly of PadlockSeq and TritSeq combinatorial barcode libraries. **A-E.** Assembly data for the PadlockSeq combinatorial barcode library are shown at left and for the TritSeq combinatorial barcode library at right. **A.** Agarose gel electrophoresis analysis of MCS size across site-wise combinatorial barcode assembly. Restriction digestion of AP plasmid libraries at each stage of barcode assembly using Transfer Sites A and B (MreI and MauBI). MCS fragments are indicated with arrowheads (**Methods**). **B.** Density plot of MCS lengths extracted from whole plasmid reads from the same AP libraries shown in A. MCS length densities are color-coded by AP library (n reads: AP_Kan_: 3, 456; PadlockSeq: Site 1, 146; Site 2, 39, 637; Site 3, 434, 845; TritSeq: AP_Amp_: 208, 873; Site 1, 125, 468; Site 2, 103, 332; Site 3, 196, 736; Site 4, 93, 934). **C.** Estimates of the number of experimentally recovered barcoded plasmids versus theoretical barcode diversity across site-wise assembly using colony-forming unit (CFU) calculations (**Methods**). Grey dots indicate the theoretical barcode diversity expected at each site. Green dots indicate the estimated number of harvested *E. coli* transformants with barcoded plasmid. The estimated fold-coverage of barcode diversity at each stage of assembly and subsequent transfer of the fully assembled combinatorial barcode into the AP with orthogonal antibiotic resistance is shown in magenta. **D.** Rank plots of the barcode element percentage abundances in the completed PadlockSeq and TritSeq libraries. In the PadlockSeq library, each set of 256 barcodes is color-coded by site. In the TritSeq library, each of the three barcode identities is color-coded and position identity is indicated by shape (A, circle; B, square; C, diamond; D, triangle). Site 4, position A barcodes 2 and 3 are indicated. Gray band indicates 95% confidence intervals expected under uniform barcode abundance (**Methods**). **E.** Sankey plots showing the relative abundances of barcode sequences at each site in the completed PadlockSeq and TritSeq combinatorial barcode libraries (n reads: PadlockSeq, 277, 401; TritSeq, 50, 336).

**Figure 9.**
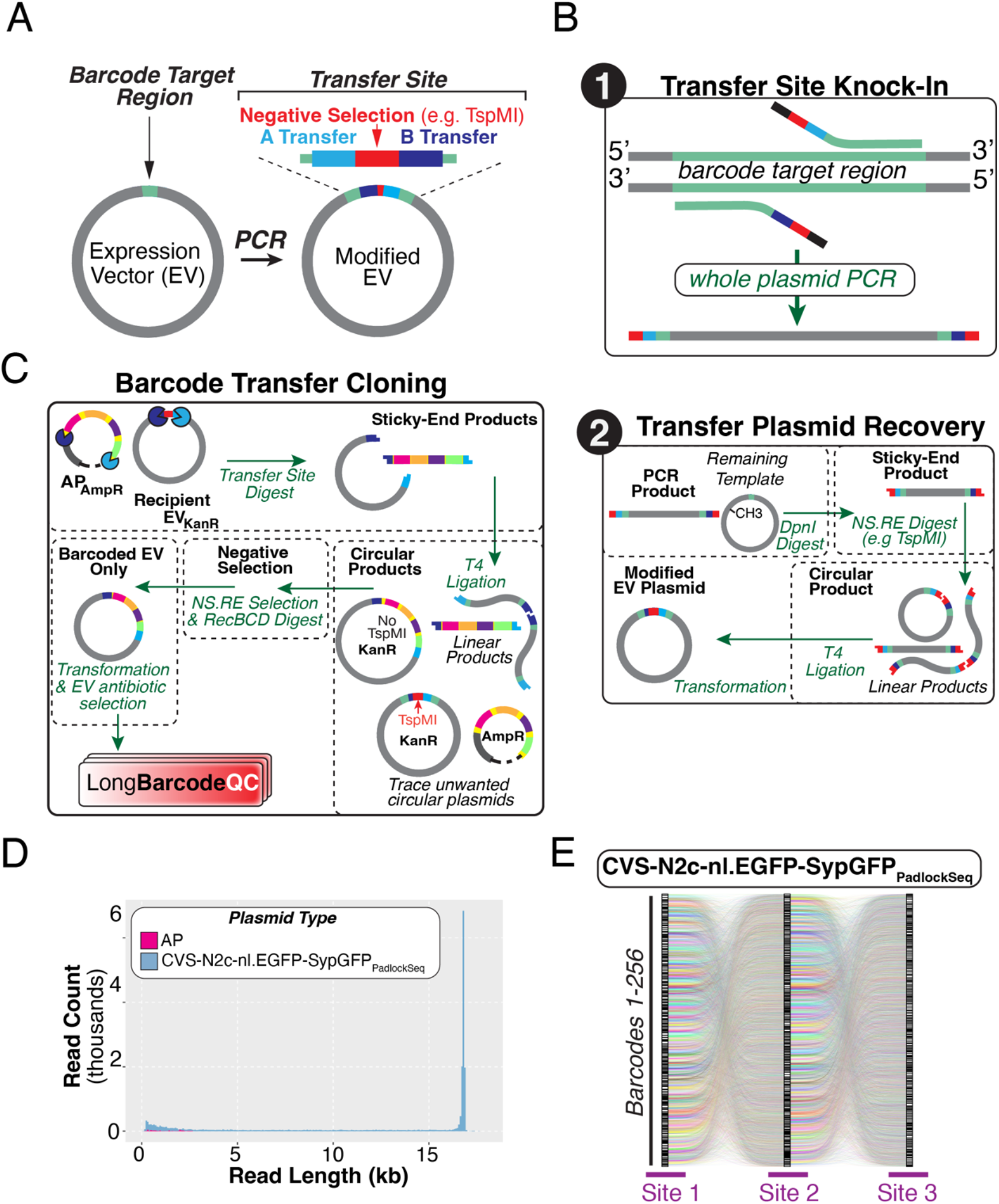
Transfer of completed combinatorial barcodes from the Assembly Plasmid to a modified recipient Expression Vector. **A.** Overview schematic of the targeted modification introduced in an Expression Vector (EV) to facilitate transfer cloning of combinatorial barcodes from an AP library. Whole-plasmid PCR introduces a Transfer Cassette comprising 8-bp Transfer Sites A and B (light and dark blue; see Figure 2B) flanking a user-selected restriction site (red) used for ligation of the modified EV and negative selection after transfer barcode cloning (e.g. TspMI). **B.** Detailed schematic of EV modification. In Step 1, forward and reverse PCR primers with homology to the target region (teal) and Transfer Site elements in their tails are used for whole plasmid PCR. In Step 2, the linear double-stranded DNA from the PCR reaction processed to generate the modified EV through the following steps: removal of residual methylated template plasmid by DpnI digestion; generation of sticky ends using the negative-selection restriction enzyme (NS.RE; e.g., TspMI); ligation with T4 DNA ligase; and transformation into *E. coli* for plasmid preparation. Dotted boxes delineate DNA cleaning or transformation steps (**Methods**). **C.** Detailed schematic of combinatorial barcode transfer cloning. Digestion at Transfer Sites A and B of the modified EV and AP library generates a sticky-ended EV backbone and barcoded MCS fragment. After EV and barcode fragment ligation, NS.RE digestion linearizes modified EV plasmids with failed barcode insertions; optional RecBCD (Exonuclease V) treatment can be used to further deplete these linear DNA species. Antibiotic selection during *E. coli* transformation enables propagation of barcoded EVs while restricting propagation of residual donor AP. *LongBarcodeQC* of whole plasmid sequencing data evaluates the purity, integrity and diversity of barcoded EV preparations. **D-E.** Example quality control of PadlockSeq transfer cloning from the AP_Kan_ into the modified CVS-N2c-nl.EGFP-SypGFP using whole plasmid sequencing reads and *LongBarcodeQC* (Supplementary Data File 1) **D.** Read length histogram color-coded by read identity (n = 15, 228 reads). **E.** Sankey plot illustrating the relative abundances of barcode sequences at each of the three PadlockSeq sites in the CVS-N2c-nl.EGFP-SypGFP_PadlockSeq_ plasmid library.

### Assembly Plasmid design

The two Assembly Plasmids (APs) were engineered to enable efficient, modular construction and transfer of high-complexity combinatorial barcode libraries (**Figure 2A**). The AP backbone is intentionally minimal, consisting only of the multiple cloning site (MCS), a bacterial origin of replication, and an antibiotic resistance cassette, resulting in compact plasmids of 1.99 kb (AP_Amp_) or 1.92 kb (AP_Kan_). This streamlined architecture reduces extraneous sequence and limits the propagation of degraded plasmid species during iterative cloning.

To enable precise transfer of completed barcode cassettes into downstream expression vectors, the entire MCS region containing sites 1-6 is flanked by two pairs of eight–base pair restriction enzyme recognition sequences (“Transfer Sites”) (**Figure 2B**). Recipient vectors are minimally modified to contain matching Transfer Sites, allowing directional excision and insertion of fully assembled combinatorial barcode libraries. We generated AP variants harboring either ampicillin (AP_Amp_) or kanamycin (AP_Kan_) resistance cassettes, enabling antibiotic selection orthogonal to the recipient expression vector and reducing carryover of the AP during transfer cloning.

At the core of the AP design is a custom MCS composed of six adjacent barcode insertion sites (sites 1–6) (**Figure 2B**). Each site contains a trio of unique restriction enzyme recognition sequences arranged in a left–middle–right configuration. The left and right enzymes generate compatible sticky ends for directional ligation of double-stranded barcode fragments, while the central restriction site enables negative selection by selectively digesting plasmids lacking successful barcode incorporation. Sites 1-4 follow the original STICR architecture (5), and sites 5 and 6 extend the trio logic to expand combinatorial capacity. Adjacent sites recycle flanking restriction sequences - for example, the BamHI recognition sequence serves as the right boundary of Site 1 and the left boundary of Site 2 - facilitating iterative, site-wise barcode incorporation.

**Figure 2C** depicts the stepwise cloning workflow, including restriction digestion, barcode ligation, negative selection, and transformation, with representative examples of sequential barcode incorporation at sites 1 and 2. To ensure assembly fidelity and quantify barcode diversity at each stage, we developed *LongBarcodeQC*, a long-read sequencing–based analytical framework designed by default to operate on the AP architecture and described in detail below. Together, these AP features support accurate, scalable, and quality-controlled assembly of combinatorial barcode libraries by avoiding suboptimal or erroneous barcode integration.

### Sequence-aware combinatorial barcode design with BarcodeEZ

To support scalable and constraint-aware barcode assembly within the AP framework, we developed *BarcodeEZ*, a Python-based software package for programmable combinatorial barcode design (**Figure 3; Methods**). *BarcodeEZ* formalizes barcode construction as a site-aware design object, allowing users to define barcode libraries corresponding to specific insertion sites within the AP architecture.

As illustrated in **Figure 3A**, users instantiate a barcode object by specifying the number of insertion sites (three sites in the example shown). Barcode sequences are then generated independently for each site by sampling from a curated corpus of orthogonal 60-mer DNA sequences that lack similarity to the transcriptome of common model organisms (**Methods**). Users can supply alternative sequence corpora to meet their needs. Following generation, sequences are subjected to validation using the `validate()`method, which removes sequences containing disallowed motifs and repopulates sites iteratively until all design constraints are satisfied. The minimal *BarcodeEZ* workflow therefore consists of object instantiation (`Barcodes()`), barcode generation (`generate_barcodes()`), and constraint-aware validation (`validate()`), followed by export of synthesis-ready oligonucleotide sequences (`write_order_form()`).

**Figure 3B** highlights additional programmable features that expand design flexibility. First, users may append fixed sequences to the left or right boundaries of barcode elements at each site, enabling incorporation of invariant motifs required for downstream applications. Second, individual sites may be subdivided into multiple internal “positions, ” each independently populated from the sequence corpus (`add_positions()`). *BarcodeEZ* automatically assigns selective sticky-end overhangs to position-specific fragments to preserve ordered restriction cloning within the AP based on a curated set of 4-mers previously benchmarked for high-efficiency directional cloning (36). The structural visualization method `show_structure()` allows inspection of site and position architecture prior to synthesis.

Importantly, *BarcodeEZ* validation is context-aware with respect to the AP cloning framework. As shown in **Figure 3C**, each candidate barcode sequence is evaluated not only in isolation but also after *in silico* incorporation into the AP restriction-site context. The `validate()` method excludes sequences that introduce unwanted restriction sites required for cloning or transfer, as well as user-defined motifs, ensuring compatibility with iterative site-wise assembly. Sequences failing validation are replaced with new sequences sampled from the corpus.

Finally, *BarcodeEZ* exports site-and position-specific barcode fragments with appropriate flanking sequences and overhangs in formats directly compatible with commercial oligonucleotide pool synthesis. This functionality bridges *in silico* design and molecular assembly, supporting accurate generation of high-complexity combinatorial barcode libraries across diverse architectures.

### Design and validation of distinct combinatorial barcode architectures

To demonstrate the generalizability of the AP and *BarcodeEZ* framework, we designed two combinatorial barcode architectures: PadlockSeq and TritSeq (**Figure 4** and **Figure 5**). Both designs encode millions of unique combinatorial barcode sequences, but differ in their structural organization and probe-based decoding logic.

We first designed a TritSeq architecture geared toward multiplexed fluorescent *in situ* hybridization (**Figure 4A**). In this configuration, each of four AP sites (1–4) is further subdivided into internal “positions” (A-D), each of which is populated with one of three possible 30 base pair (bp) barcode sequences. This three-state encoding scheme is well suited for decoding using spectrally distinct fluorescent *in situ* hybridization probes (**Figure 4B**). The 16 site/position design efficiently expands combinatorial barcode diversity 3^16^ (n = 43, 046, 721 total), with 3 x 16 (n = 48) probes. In this study, the TritSeq design is presented to highlight the flexibility of the AP and *BarcodeEZ* design and assembly framework to implement site-internal positions as a means for highly efficient probe-based combinatorial barcode decoding.

The PadlockSeq architecture distributes combinatorial diversity across three independent insertion sites (sites 1–3) within the AP (**Figure 5A**). At each site, a 25 bp site-variable region is flanked by site-conserved sequences that support padlock probe hybridization and binding of a site-specific primer. These conserved elements enable padlock probe circularization followed by rolling-circle amplification initiated by the site-targeted primer, generating localized amplification of barcode-containing transcripts (**Figure 5B**)(18, 27). In our case, 256 distinct site-variable sequences were designed for each of the three sites, enabling 256^3^ (n = 16, 777, 216 total) combinatorial barcode identities to be decoded with 256 x 3 (n = 768) padlock-primer probe pairs.

As illustrated in **Figure 5B**, each padlock probe incorporates a 5-bp internal sequencing identifier decoded by *in situ* sequencing. The first four bases encode the barcode number (1–256), while the fifth base specifies the site location (sites 1–3), allowing both barcode element number and the site location to be determined in a single sequencing read. Thus 75% of the 4^5^ (n = 1, 024) total probe sequence identifiers are used to determine barcode identity, leaving the remaining 25% (n = 256) available for additional experimental needs, such detecting gene-specific transcripts that delineate cell type or state populations. While each barcoded mRNA contains all three sites, site-specific amplification is in practice mutually exclusive, thus combinatorial barcode identities are reconstructed by aggregating site-resolved sequencing calls across sites 1–3 within each cell.

To validate *in vivo* detection of PadlockSeq barcodes, we generated six CVS-N2c rabies virus (RABV) genomes encoding barcodes with distinct site-variable sequences at each site (A-F)(**Figure 5C**). Barcode sequences were inserted into the 3’ UTR of an NLS-GFP reporter gene, replacing the native glycoprotein gene, a spread-incompetent configuration that leads to barcode expression in only directly transduced cells. Viruses were rescued with the native N2c glycoprotein envelope, mixed at equal titers, and injected intracerebrally as a cocktail into the mouse primary somatosensory cortex (**Methods**). Barcode readout was performed using padlock probe-based *in situ* sequencing on Pyxa (Stellaromics), which implements a version of STARmap optimized for thick tissue sections (18, 27). Following padlock probe hybridization and rolling-circle amplification, 5 bp probe identifiers were decoded from a set of 264 neurons found in a single field of view around the injection site (**Figure 5D**; **Methods**).

Sequenced site-specific amplicons (“decodes”) amplified from barcoded mRNAs enabled sensitive and accurate reconstruction of complete PadlockSeq combinatorial barcode identities in each neuron (**Figure 5D, E; Methods**). Decodes for each site were reliably recovered (mean decode count per cell: Site 1, 45.8±1.9 (s.e.m); Site 2, 20.7±1.2; Site 3, 25.1±1.5), identifying both single and multiple infection events in individual neurons. These decoding results validate that the PadlockSeq barcode architecture is well-suited for tracking millions of distinct RABV-expressed RNAs in infected neurons in intact brain tissue sections using probe-based *in situ* sequencing on the Pyxa platform.

### LongBarcodeQC supports highly accurate combinatorial barcode assembly through analysis of whole plasmid nanopore sequencing

To support rapid and cost-effective quality control of combinatorial barcode libraries during serial AP cloning, we developed *LongBarcodeQC*, a Python-based command-line analysis package for whole plasmid nanopore sequencing data (**Figure 6**; **Methods**). *LongBarcodeQC* evaluates barcode insertion structures within the MCS, restriction site integrity, and barcode representation directly from long-read FASTQ files generated at intermediate or final stages of cloning.

As shown in **Figure 6A**, the minimal *LongBarcodeQC* workflow requires four inputs: 1) nanopore sequencing reads (FASTQ), 2) a reference plasmid sequence (FASTA; default: AP), and 3) user-defined barcode sequences (FASTA) exported from *BarcodeEZ* and 4) expected insert lengths. Optional parameters allow specification of MCS flanking sequences, restriction sites used for cloning and negative selection, expected insertion numbers, and toggling of AP-specific metrics. Following adaptor trimming, reads are aligned to both the reference plasmid and the *E. coli* genome to distinguish full-length plasmid molecules from bacterial genomic contamination. Plasmid-aligned reads are oriented to a common strand and the MCS region is extracted using defined flanking anchor sequences. Barcode sequences are then aligned to each extracted MCS and barcode identities are assigned based on alignment score thresholds. *LongBarcodeQC* generates an interactive HTML report summarizing results, along with a detailed CSV file containing per-read metadata (**Methods**).

The summary HTML output integrates four primary analyses to evaluate site-specific barcode cloning outcomes (**Figure 6B–E**). First, alignment summaries stratify read-length distributions by alignment category (reference plasmid-aligned, *E. coli*-aligned, or unaligned reads), enabling rapid assessment of plasmid enrichment and sequencing quality (**Figure 6B**). Second, MCS length distributions are extracted at base-pair resolution from plasmid-aligned reads, and barcode assignments are overlaid to quantify expected insertion sizes and detect aberrant cloning products (**Figure 6C**). Third, an interactive restriction site analysis reports the presence or absence of expected restriction sequences within each extracted MCS, facilitating confirmation of successful site-specific cloning and identification of failures in negative selection steps (**Figure 6D**). Fourth, barcode calling metrics are derived from alignment scores of user-specified barcode sequences to each MCS. A Z-score distribution of barcode alignment scores enables visualization and thresholding for confident barcode assignment (**Figure 6E**).

Together, *LongBarcodeQC* provides a plasmid-and barcode-aware quality control framework that integrates seamlessly with *BarcodeEZ* outputs and the AP structure. By leveraging whole plasmid nanopore reads, *LongBarcodeQC* resolves complete MCS assemblies at base-pair resolution from single molecules, enabling sequence-resolved evaluation of barcode insertion during site-wise barcode assembly. Because nanopore sequencing can be performed rapidly and at low per-sample cost, this approach supports efficient optimization and quality control of combinatorial barcode library construction.

### Validation of site-wise serial assembly quality control using LongBarcodeQC

To demonstrate how *LongBarcodeQC* analysis informs experimental optimization during iterative barcode assembly, we analyzed PadlockSeq barcode cloning into site 2 using the BamHI–KpnI–NheI restriction enzyme trio (**Figure 7A**). Following AP digestion with BamHI and NheI, barcode fragments bearing compatible overhangs were ligated into site 2. The native site 2 region spans 24 bp; insertion of a single-site PadlockSeq barcode yields a modified 62 bp site 2, whereas multiple tandem insertions produce larger products (∼174 bp). These defined sequence outcomes provide a quantitative basis to evaluate insertion multiplicity and restriction enzyme-based negative selection during serial cloning.

To highlight how experimental parameters such as ligation conditions influence insertion multiplicity, we compared plasmid libraries generated using AP:barcode fragment molar ratios of 1:1.5 and 1:0.3. *LongBarcodeQC* analysis of extracted MCS regions revealed a dominant peak at 247 bp, corresponding to the expected MCS length with a single barcode insertion. Reducing the ratio of barcode fragment to AP vector decreased the percentage of MCS sequences with expected lengths of multiple insertions (+112 bp) from 5.7% to 2.5% (**Figure 7B**). These quantitative results illustrate how *LongBarcodeQC* facilitates optimization of assembly conditions against user-defined benchmarks for acceptable multi-insert frequencies.

In addition to expected single and multiple insert MCS populations, a minor class of reads encoded MCS sequences 17 bp shorter on average than predicted for single barcode insertions (**Figure 7B**). Careful inspection of AP alignments for this population of reads revealed short deletions throughout the MCS, consistent with known nanopore sequencing error profiles rather than cloning artifacts (42) (**Figure 7C**). Such results highlight how the *LongBarcodeQC* framework helps distinguish bonafide barcode insertion heterogeneity from nanopore sequencing errors.

We next evaluated the efficacy of restriction enzyme-based negative selection by comparing MCS lengths and sequence properties from sequential site 2 and site 3 barcode fragment cloning experiments. In correctly assembled constructs, insertion of a site 2 barcode disrupts the internal KpnI recognition sequence, whereas empty vectors of failed ligations retain this site (**Figure 7A**). Following site 2 barcode cloning, *LongBarcodeQC* analysis identified a minor population (∼10%) of reads that retained the KpnI site, lacked site 2 barcode calls and exhibited MCS lengths consistent with failed insertion (**Figure 7D**). Following KpnI digestion upstream of subsequent site 3 barcode cloning, plasmids with site 2 cloning failures were removed from the site1-2-3 plasmid library (< 0.04 %). The resulting library displayed MCS length distributions and barcode-calling patterns consistent with successful barcode incorporation at sites 1-3.

Collectively, these examples demonstrate how AP design features coupled with whole plasmid nanopore sequencing and *LongBarcodeQC* support both experimental optimization and experiment-to-experiment quality control. Thus, the AP/*LongBarcodeQC* framework helps users achieve single-insertion assembly benchmarks during the serial assembly of complex combinatorial barcode libraries. Having established optimized ligation conditions and validated restriction enzyme-based negative selection, we applied this framework to construct complete PadlockSeq and TritSeq combinatorial libraries.

### Construction of PadlockSeq and TritSeq combinatorial barcode libraries

We constructed full PadlockSeq and TritSeq combinatorial barcode libraries in the AP through serial, site-specific restriction cloning and negative selection, evaluating each assembly stage with whole plasmid nanopore sequencing and *LongBarcodeQC* analysis (**Figure 8**; **Methods**). Barcode libraries were assembled in either the ampicillin-resistant AP backbone (AP_Amp_) or the kanamycin-resistant backbone (AP_Kan_) and subsequently transferred as intact, fully barcoded MCS cassettes into the orthogonal AP backbone to facilitate downstream transfer into diverse types of expression vectors. The complete PadlockSeq and TritSeq AP plasmid libraries have been deposited with Addgene to enable community access.

For the PadlockSeq design (3 sites, one position per site, 256 sequences per site; ∼16.8 million combinations) and the TritSeq design (4 sites, four internal positions per site, three sequences per position; ∼43 million combinations), stepwise insertion of barcode fragments produced the predicted incremental increases in MCS length (**Figure 8A, B**). Agarose gel electrophoresis revealed the expected discrete size shifts in the MCS corresponding to successful incorporation at each site. *LongBarcodeQC* analysis confirmed the expected MCS length distributions at base pair resolution.

To preserve barcode diversity during transformation and plasmid propagation, we quantitatively benchmarked library coverage at each stage of site-wise assembly. For intermediate and final plasmid libraries, theoretical barcode diversity was compared to empirical estimates of barcode-containing colony-forming units (CFUs) recovered from large-format plate-based transformations (**Figure 8C**; **Methods**). Although we targeted >5-fold coverage of theoretical library diversity at each assembly stage, estimated coverage of the final PadlockSeq and TritSeq libraries was 9.9-fold and 1.9-fold, respectively.

We assessed the efficacy and completeness of PadlockSeq and TritSeq assembly using whole plasmid long-read sequencing and *LongBarcodeQC* outputs. Specifically, we assessed barcode diversity at each site/position followed by the degree of independent assortment of these elements into full combinatorial barcodes (**Figure 8D, E**). To evaluate site/position barcode diversity, we used Simpson evenness (*E_2_*) as a metric. *E_2_* is normalized to a maximum of 1, with higher values indicating more uniform barcode representation (**Methods**). PadlockSeq *E_2_* values ranged from 0.83-0.89 across sites 1-3, while TritSeq *E_2_* ranged from 0.93-0.99 except for site 4, position A (0.63). These results suggest that the diversity of site/position barcodes was high for both libraries, with one exception for TritSeq. To evaluate randomness of combinatorial barcode assembly, we used permutation-adjusted normalized mutual information (NMI) to quantify statistical dependence of site/position barcodes, accounting for the finite-sampling bias of our measurements (**Methods**). NMI values close to 0 indicate independent assortment, whereas increasingly positive values indicate progressively stronger non-random associations between barcode identities at different sites/positions. NMI values were near zero for both libraries, indicating highly independent barcode assortment. Mean NMI was 0.0022 for PadlockSeq (maximum, 0.0035) and 0.0004 for TritSeq across 120 site/position pairs (maximum, 0.0107). These results indicate minimal dependence between barcode identities at different sites in either library. At the level of complete combinatorial barcodes, the estimated probability that two randomly sampled molecules carried the same complete combinatorial barcode was approximately 1 in 1.6 million for PadlockSeq and 1 in 5.5 million for TritSeq (**Methods**).

### Expression vector transfer of fully assembled combinatorial barcodes

To enable flexible experimental use of combinatorial barcode libraries, we developed a modular strategy to modify expression vectors (EVs) for barcode transfer cloning. Target EVs are engineered to contain a Transfer Cassette comprising two 8-bp restriction sites (Transfer Sites A and B) flanking an internal restriction site used for negative selection (**Figure 9A**). These Transfer Sites match those flanking the MCS in the AP. Importantly, orthogonal antibiotic resistances between AP and the EV further prevent AP carryover during barcode transfer cloning.

The modified EV is generated by whole plasmid PCR using primers encoding the Transfer Cassette, followed by a molecular workflow to regenerate a clean circular plasmid (**Figure 9B**; **Methods**): template EV plasmid is removed by DpnI digestion; the negative selection site is digested from the terminal ends of the linear PCR product, generating overhangs suitable for ligation, and residual linear DNA species are degraded enzymatically. Modified EV suitable for barcode transfer cloning is circularized through ligation, transformed, and recovered (**Methods**).

For transfer cloning (**Figure 9C**), the fully assembled barcode cassette is excised from the AP by digesting Transfer Sites A and B and ligated into the modified EV backbone after equivalent Transfer Site digestion. Restriction enzyme–based negative selection removes EV plasmids with failed barcode ligation, while antibiotic selection eliminates residual AP backbone after transformation. This procedure selectively recovers plasmid preparations that contain only barcoded EV.

We applied this strategy to transfer the completed PadlockSeq library from AP_Kan_ into the modified 16.7 kb CVS-N2c rabies virus genome plasmid used for *in vivo* validation (**Figure 9D**; **Methods**). Whole plasmid nanopore sequencing coupled with *LongBarcodeQC* confirmed correct cassette insertion within the full-length viral genome backbone, with minimal AP carryover and expected MCS architecture (**Figure 9D, E**).

This transfer cloning procedure demonstrates how complex combinatorial barcode libraries accurately assembled in a streamlined, purpose-built AP can be transferred to a large and unwieldy expression vector with minimal loss of barcode diversity or generation of unwanted byproducts, enabling high-quality plasmid preparations suitable for downstream probe-based genomics applications.

## DISCUSSION

Probe-based genomics methods are extending single-cell analysis into intact tissue and fixed-cell contexts, yet reagents that facilitate highly multiplexed experimentation on these platforms remain limited. Combinatorial barcoding – a strategy that uses combinations of known synthetic sequences to generate an exponentially larger set of unique molecules - offers a powerful general strategy for highly multiplexed experimentation: specific combinatorial barcodes can be linked to particular experimental conditions and then decoded downstream using a limited set of sequence-targeting probes (28, 33–35). However, generating highly accurate combinatorial barcode libraries designed to fit the needs of existing and future probe-based genomics technologies remains a challenge. Despite decreased cost and improved scalability, *de novo* synthesis of libraries containing millions of combinatorial barcodes long enough to accommodate multiple probe target sites remains prohibitively expensive. As a reference point, at current synthesis costs of $0.01-0.001/bp, the PadlockSeq and TritSeq libraries we design and assemble in this study would cost 2.7-27 million and 23.8-238 million dollars to synthesize *de novo*, respectively. Thus, methods that enable design and accurate assembly of structurally diverse combinatorial barcode architectures remain an outstanding need.

To address this need, we developed an integrated design–assemble–validate–deploy framework for combinatorial DNA barcodes tailored to probe-based readouts. This framework integrates a streamlined Assembly Plasmid (AP), computationally aided barcode design through *BarcodeEZ*, long-read structural validation using nanopore sequencing and *LongBarcodeQC* analysis, and modular barcode transfer into expression vectors of interest. We demonstrate that our integrated framework offers flexible and assembly-aware combinatorial barcode design; confirm near-random assembly of diversely barcoded elements; and validate that our strategy helps avoid assembly errors and unwanted byproducts that can arise during serial restriction cloning.

A central goal of this work was flexible and precise engineering of complex barcode architectures. *BarcodeEZ* formalizes barcode design as a site-aware process that operates within the constraints of the AP sequence structure. Barcode sequences are sourced from pre-defined corpuses, allowing users to define barcode libraries with synthetic sequences orthogonal to model species transcriptomes or other constraints. *BarcodeEZ* enables incorporation of site-specific fixed sequences, subdivision of AP sites into internal barcode positions, and enforcement of sequence constraints for seamless AP cloning. The flexibility of *BarcodeEZ* and design integration with the AP enables users to develop custom combinatorial barcode libraries optimized for detection using distinct probe-based decoding strategies, exemplified by the distinct PadlockSeq and TritSeq combinatorial barcode libraries we assembled and make publicly available.

We sought to minimize the introduction and propagation of unwanted cloning products, a major challenge of site-wise barcode assembly. The AP was intentionally engineered as a compact plasmid lacking extraneous sequence, minimizing the truncation and persistence of plasmid species that retain antibiotic resistance. We implemented this streamlined AP design after our previous attempts to assemble combinatorial barcodes in larger vectors consistently resulted in plasmid contaminants that outcompeted our target vectors after transformation in *E. coli*. We also previously observed AP carryover during barcode transfer cloning into expression vectors when antibiotic resistance genes were shared; we thus implemented two AP variants with distinct antibiotic resistance cassettes to enable orthogonal antibiotic selection relative to the recipient expression vector. Extending the restriction site “trio” logic developed by STICR (5), we validated that restriction enzyme-based negative selection effectively removes plasmids lacking successful barcode incorporation at each stage of assembly. Taken together, enzymatic negative selection, per-site sequencing-based quality control, and AP/expression vector antibiotic orthogonality, provide complementary mechanisms to suppress insertion failures and plasmid byproducts that are otherwise difficult to eliminate by size selection alone. Importantly, mutations in the AP backbone and multiple barcode insertions at single sites cannot be removed enzymatically and therefore must be quantitatively assessed after each stage of assembly through sequencing.

Whole plasmid nanopore sequencing integrated with *LongBarcodeQC* analysis underpins our quality-control strategy during combinatorial barcode assembly. Because the AP backbone and site-specific barcode sequences are known *a priori*, long-read sequencing resolves complete multiple cloning site (MCS) structures at base pair resolution from individual molecules in a manner that is robust and distinguishable from nanopore sequencing errors. In practice, low-coverage whole plasmid nanopore sequencing is a rapid and affordable strategy sufficient for quality control. While we generated the long-read sequencing data in our laboratory using low-throughput “flongle” flowcells (Oxford Nanopore Technologies) in just a few hours, similar whole-plasmid sequencing datasets can be acquired from third-party commercial services with turnaround times of less than 2 days. Of note, short-read sequencing is significantly more expensive, typically requires days-long runtimes, and may be insufficient for sampling complete combinatorial barcode molecules. Thus whole plasmid long-read sequencing may be preferable for many investigators. In practice, combining read-specific analysis of MCS lengths, barcode assignments, and the presence of MCS restriction enzyme recognition sequences, proves to be a highly effective strategy for diagnosing cloning outcomes. These metrics are available as standard interactive HTML-based outputs from *LongBarcodeQC*, while verbose *LongBarcodeQC* outputs are available for custom analysis.

Separation of combinatorial barcode assembly in the AP from downstream deployment in expression vectors serves two important functions. First, as noted above, assembly in the AP drastically reduces unwanted byproducts that tend to arise during serial restriction cloning with larger vectors and facilitates accurate propagation and sharing of completed libraries. Second, by developing a transfer strategy using a restriction enzyme-based trio targeted to the expression vector of interest, AP-based combinatorial barcodes can be modularly transferred to diverse expression vectors to support different experimental needs. In our study, we showcase transfer cloning of the completed PadlockSeq library into a16.7 kb CVS-N2c rabies virus genome plasmid. Our results demonstrate how this modification and transfer process is compatible with large and complex expression vectors, maintaining barcode representation and avoiding contamination by the AP or parental expression vector. Platform-specific use of the same combinatorial barcodes delivered by distinct expression vectors narrows optimization to probe delivery conditions rather than wholly new barcode schemes and probe sets.

The sensitivity of barcode detection depends on both the properties of the expression vector and the characteristics of the probe-based readout platform. We validated PadlockSeq combinatorial barcode decoding *in vivo* following rabies virus transduction of mouse cortex using STARmap *in situ* sequencing on the commercial Pyxa platform (18, 27). These results support the PadlockSeq architecture as a probe-efficient strategy for decoding millions of RNA-linked experimental manipulations with high fidelity. Interestingly, these validation experiments suggest that unique RNA barcodes delivered into individual cells may also help delineate cell boundaries. This is important because accurate cell segmentation in complex tissue remains a major challenge for spatial transcriptomic data (43–46). Future experiments that use probe sets to detect PadlockSeq RNA barcodes alongside host cell RNAs should enable more accurate single-cell molecular profiling than host cell RNAs alone, especially if fluorescent proteins are used to image cellular morphology upstream of RNA analysis (27). Thus PadlockSeq RNA barcodes decoded on the Pyxa platform could facilitate scaling CRISPR-based cell morphology screens from a handful of genes to the full genome (47), enable high-throughput reconstruction of synaptic networks in the intact brain(13), or support large-scale screening of engineered fluorescent reporter variants for improved monitoring of cell signaling pathways (48). Following the PadlockSeq model, we recommend that new combinatorial barcode designs are platform-validated using a handful of known combinatorial barcodes before full libraries are assembled.

Several limitations merit consideration. Improvements in the cost and scalability of long synthetic DNA constructs could reduce the need for serial restriction-based combinatorial barcode assembly. However, even if DNA synthesis costs plummet, the assembly-aware design, structural validation, and modular transfer strategies we introduce remain important even for combinatorial barcode libraries generated by alternative methods. The TritSeq design was presented to highlight the generalizability of our combinatorial barcode design and assembly approach, including the use of site-internal positions to extend the number of combinatorial barcode sites beyond the six restriction site trios engineered into the AP. We did not validate TritSeq barcode detectability in our study. As is the case for all barcode schemes, sensitivity of detection depends on properties of the expression vector, tissue context, and the experimental approach, as well as the probe-based readout platform. While we validated sensitive and accurate detection of PadlockSeq combinatorial barcode architecture in the mouse brain using six known examples, detection of the full PadlockSeq repertoire using 768 probes could in theory be more variable. In our study, we used conservative estimates of barcode plasmid-containing colony-forming units to guard against bottlenecking barcode diversity during site-wise assembly of the PadlockSeq and TritSeq libraries. However, representation of individual PadlockSeq and TritSeq combinatorial barcodes deviated somewhat from uniformity. After carefully avoiding bottlenecking barcode diversity during transformation of *E. coli*, our sequencing data indicate that biases in the representation and ligation of oPool sequences at each site/position have relatively minor effects, whereas biased replication in *E. coli* may have a larger effect. For example, we speculate such biased replication was responsible for the trade-off in representation between barcode sequences 2 and 3 at Site 4 Position A in the TritSeq library, but other mechanisms could be responsible. The greater length of combinatorial barcodes may make them more susceptible to these cryptic replication effects than shorter barcode sequences. Lastly, sequencing the PadlockSeq and TritSeq libraries to saturation using alternative single-molecule sequencing technologies could complement our estimates of effective barcode diversity.

In summary, we describe a modular and quality-controlled framework for constructing probe-compatible combinatorial barcode libraries. By integrating programmable design, serial restriction-based assembly, long-read structural validation, and flexible transfer into expression vectors, this platform fills a technically demanding niche by providing precise engineering and straightforward deployment of combinatorial barcode reagents. As probe-based genomics technologies continue to evolve, standardized and shareable combinatorial barcode libraries stand to function as critical reagents to support highly multiplexed experimentation in intact tissues and fixed cells.

## DATA AVAILABILITY

The Assembly Plasmids, TritSeq, and PadlockSeq plasmid libraries described in this article will be available from Addgene upon peer-reviewed publication (Addgene ID numbers: *Plasmids*: AP-Kan, 261892; AP-Amp, 261893. *Pooled Libraries*: PadlockSeq(AP-Amp), 261888; PadlockSeq(AP-Kan), 261889; TritSeq(AP-Amp), 261890; TritSeq(AP-Kan), 261891). Prior to Addgene distribution, reagent requests should be directed to Dr. Saunders. Source code, documentation, and installation instructions for *BarcodeEZ* and *LongBarcodeQC* are available through GitHub at https://github.com/ArpiarSaundersLab/BarcodeEZ and https://github.com/ArpiarSaundersLab/LongBarcodeQC, respectively. Both packages are distributed as pip-installable Python software. All custom code used to generate the analyses and figures presented in this manuscript is available at https://github.com/ArpiarSaundersLab/2026_combinatorial_barcode_analysis. Raw Oxford Nanopore whole plasmid sequencing data generated in this study will be deposited in the NCBI Sequence Read Archive (SRA) before publication. Processed *LongBarcodeQC* outputs and associated summary data are provided in Supplementary Data File 5.

## SUPPLEMENTARY DATA

Supplementary Data File 1 - Plasmid maps (.dna)

Supplementary Data File 2 - PadlockSeq barcode sequences (.csv)

Supplementary Data File 3 - TritSeq barcode sequences (.csv)

Supplementary Data File 4 - PadlockSeq probes and site-conserved primers (.csv)

Supplementary Data File 5 - *LongBarcodeQC* processed datasets and HTML summary files

Supplementary Data Files are available at https://github.com/ArpiarSaundersLab/2026_combinatorial_barcode_analysis

## AUTHOR CONTRIBUTIONS

Conceptualization: Z.G., E.T., A.S.

Methodology: Z.G, E.T., L.B. K.P., K.Y., M.S., A.N., Y.F., A.S.

Software: Z.G., K.P., K.Y., J.H.

Validation: Z.G., E.T., J.Z., H.B.

Formal analysis: Z.G., K.Y., K.P., A.S.

Investigation: E.T., L.BA., M.S., A.N., J.Z., H.B

Resources: J.Z., Y.F., H.B.

Data curation: Z.G., J.H.

Writing – original draft: Z.G., E.T., A.S.

Writing – review & editing: All authors

Visualization: Z.G., E.T., A.S.

Supervision: A.S.

Project administration: A.S.

Funding acquisition: A.S.

## ACKNOWLEDGEMENTS

The authors thank Ryan Delgado and Thomas Nowakowski for sharing the original STICR plasmid (Addgene Accession #186334). The authors also thank members of the Saunders, Andrew Adey and Brian O’Roak labs at OHSU for helpful conversations, as well as Dr. Fenna Krienen (Princeton University) and Dr. Michael Ratz (Karolinska Institute) for thoughtful feedback on the manuscript.

## DECLARATION OF GENERATIVE AI AND AI-ASSISTED TECHNOLOGIES

During the preparation of this work, the authors used ChatGPT(v5.4) and Claude Opus 5 for building syntactically correct code, debugging errors, and drafting analysis code and descriptive text. ChatGPT(v5.4) was also used for editing and proofreading of manuscript text. After using these tools, the authors reviewed and edited the content and take full responsibility for all text.

